# hu.MAP3.0: Atlas of human protein complexes by integration of > 25,000 proteomic experiments

**DOI:** 10.1101/2024.10.11.617930

**Authors:** Samantha N. Fischer, Erin R. Claussen, Savvas Kourtis, Sara Sdelci, Sandra Orchard, Henning Hermjakob, Georg Kustatscher, Kevin Drew

**Affiliations:** Department of Biological Sciences, University of Illinois at Chicago, Chicago, IL 60607; Centre for Genomic Regulation (CRG), The Barcelona Institute of Science and Technology, Barcelona, Spain; European Molecular Biology Laboratory, European Bioinformatics Institute (EMBL-EBI), Wellcome Genome Campus, Hinxton, Cambridge CB10 1SD, UK; Wellcome Centre for Cell Biology, University of Edinburgh, Edinburgh EH9 3BF, UK

## Abstract

Macromolecular protein complexes carry out most functions in the cell including essential functions required for cell survival. Unfortunately, we lack the subunit composition for all human protein complexes. To address this gap we integrated >25,000 mass spectrometry experiments using a machine learning approach to identify > 15,000 human protein complexes. We show our map of protein complexes is highly accurate and more comprehensive than previous maps, placing ∼75% of human proteins into their physical contexts. We globally characterize our complexes using protein co-variation data (ProteomeHD.2) and identify co-varying complexes suggesting common functional associations. Our map also generates testable functional hypotheses for 472 uncharacterized proteins which we support using AlphaFold modeling.

Additionally, we use AlphaFold modeling to identify 511 mutually exclusive protein pairs in hu.MAP3.0 complexes suggesting complexes serve different functional roles depending on their subunit composition. We identify expression as the primary way cells and organisms relieve the conflict of mutually exclusive subunits. Finally, we import our complexes to EMBL-EBI’s Complex Portal (https://www.ebi.ac.uk/complexportal/home) as well as provide complexes through our hu.MAP3.0 web interface (https://humap3.proteincomplexes.org/). We expect our resource to be highly impactful to the broader research community.

## Introduction

Proteins are the functional units of the cell yet they carry out cellular functions by self assembling into large macromolecular protein complexes. A more complete understanding of human cells requires a better understanding of the functional complexes in those cells. Major advances in high throughput proteomics and machine learning workflows have allowed both the collection of large datasets and the integration of those datasets into more accurate and complete sets of protein complexes. In particular, experimental workflows based on affinity purification mass spectrometry (AP-MS)^1–7^, co-fractionation mass spectrometry (CF-MS)^8–13^, and proximity labeling^14,15^ have been applied to different cells, tissues, and organisms generating a large compendium of proteomics data focused on identifying protein complexes. We have previously integrated several of these datasets using machine learning in our hu.MAP1 and hu.MAP2 resources which have been extremely valuable at identifying candidate disease genes, functionally annotating the proteome’s least characterized proteins, and identifying multifunctional promiscuous proteins^16,17^. In total hu.MAP2 placed half of the human proteins (∼10,000) into protein complexes^16^.

Orthogonal approaches, based on co-expression/covariation of proteins in mass spectrometry based datasets across many conditions, have been developed to identify functional modules of the cell^18–20^. While these networks are not physical in nature, the efforts have shown subunits of protein complexes are often highly co-expressed and provide a powerful signature for shared co-complex interactions.

Here we describe an improved hu.MAP3.0 computational workflow for the integration of high throughput proteomics experiments to identify human protein complexes. Using this workflow we integrate >25,000 mass spectrometry experiments from multiple approaches (i.e. AP-MS, CF-MS, proximity labeling) and identify greater than 15,000 protein complexes for ∼70% of the human proteome (>13,700 total proteins). We compare physical complexes identified by hu.MAP3.0 to protein covariation networks and find many complexes have tight co-expression patterns. Additionally, we utilize AlphaFold structural models of protein pairs to identify mutually exclusive proteins (i.e. pairs of proteins that cannot coexist in complex). These are further supported by co-variation networks. In total we identify 511 mutually exclusive pairs which point to potential regulation and functional outcomes of complexes in different cell types, tissues, and organelles. We also analyze our map to identify understudied proteins which physically associate with well annotated complexes. We place 472 understudied proteins, including several disease associated, into complexes providing testable hypotheses as to their function. Finally, we make our identified complexes available through two major resources, a hu.MAP3.0 web resource (https://humap3.proteincomplexes.org) and EMBL-EBI’s Complex Portal (https://www.ebi.ac.uk/complexportal/home)^21^.

## Results

To construct a more accurate and comprehensive set of protein complexes, we use machine learning to integrate published high throughput proteomics experiments. Each experimental dataset targets interactions from different cell types and different parts of the proteome. This therefore provides a broad sampling of human protein complexes as well as multiple lines of evidence for each protein complex. In Figure 1A, we show a schematic representation of the full machine learning pipeline to produce protein complexes. The pipeline consists of 3 major steps:

1. organization and generation of features predictive of protein interactions,
2. training of ML model to generate protein interaction network, and
3. clustering of network to identify complexes. This final step results in 15,326 complexes covering 13,769 human proteins and consisting of 159,451 total scored protein interactions. This set of complexes, which we call the hu.MAP3.0 human protein complex map, considerably increases coverage compared to previous complex maps (Fig 1B-D).

**Figure 1.**
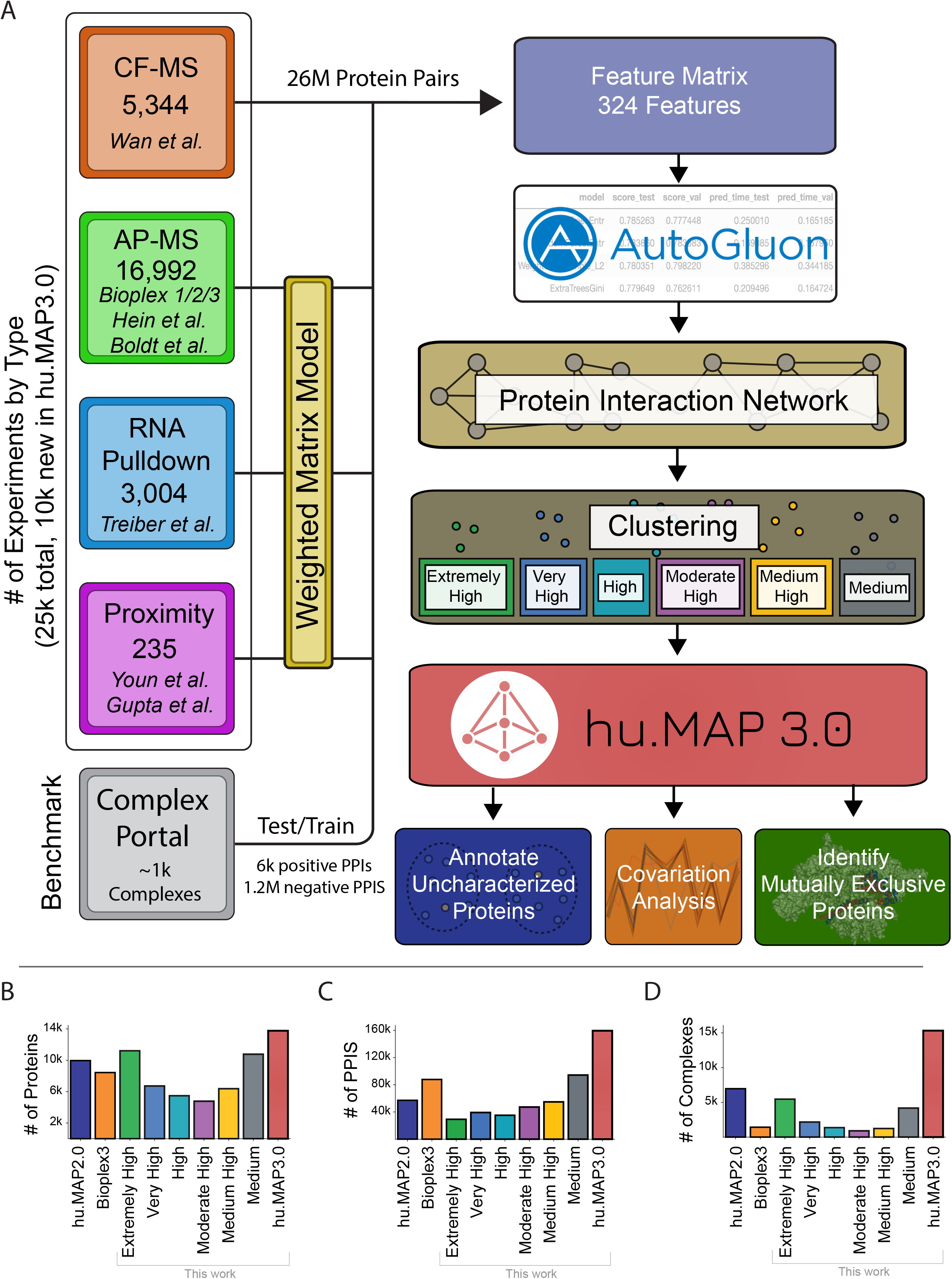
Machine learning workflow identifies human protein complexes. (**A**) Representation of the hu.MAP3.0 workflow which integrates ∼25k mass spectrometry experiments to identify protein complexes. A feature matrix for machine learning classification of protein pairs was constructed using four classes of experimental datatypes (i.e. Cofractionation Mass Spectrometry (CF-MS), Affinity Purification Mass Spectrometry (AP-MS), Weighted Matrix Model applied to RNA Pulldown Mass Spectrometry experiments, and Proximity Labeling Mass Spectrometry experiments. A reanalysis of AP-MS and proximity labeling data was performed using a Weighted Matrix Model. The feature matrix was labeled using known Complex Portal interactions and a classifier was trained using the AutoGluon model selection pipeline. A protein interaction network was built by applying the model to ∼26 million pairs of proteins that provided a confidence score of co-complex interaction. The protein interaction network was clustered producing complexes of “Extremely High” confidence to “Medium” confidence. Complexes were then analyzed to annotate uncharacterized proteins and identify mutually exclusive proteins. (**B**) Comparison of human proteome coverage between hu.MAP2.0 (dark blue), Bioplex3 (orange), hu.MAP3.0(red), and the 6 levels of complex confidence of hu.MAP3.0 (green = “Extremely High”, blue = “Very High”, aqua = “High”, purple = “Moderately High”, yellow = “Medium High”, and gray = “Medium”). The complete set of hu.MAP3.0 complexes covers 13,769 human proteins outperforming hu.MAP2.0 and Bioplex3. (**C**) Comparison of number of protein interactions (PPIs) across hu.MAP2.0, Bioplex3, hu.MAP3.0, and 6 levels of complex confidences. Same colors as in (**B**). The hu.MAP3.0 complex map considerably increases the number of total protein interactions including 159,451 total scored interactions. (**D**) Comparison of number of complexes in each dataset. Same colors as in (**B**). hu.MAP3.0 contains 15,326 complexes, a substantial increase in the number of complexes than previously published protein complex maps.

### Integration of high throughput proteomics datasets to assemble evidence of protein interactions

We first describe our development of the hu.MAP3.0 machine learning pipeline. The initial phase includes organizing evidence features from experimental datasets from CF-MS^8^, AP-MS^1–5,7^, Proximity Labeling^14,15^, and RNA pulldown experiments^22^. In our current work, we include a significant number of new experiments from the BioPlex3^1^ effort which adds ∼10k new AP-MS experiments to our workflow. Machine learning features from each experiment were collated into a feature matrix where each feature provides evidence of a protein-protein interaction and is specific to each experiment type. CF-MS features contain similarity measures of pairs of protein’s elution in a biochemical separation (e.g. Pearson correlation coefficient) with high similarity as evidence for potential interaction. Pulldown and proximity features contain metrics of the presence of an individual prey protein identified in a bait protein experiment.

These measures provide models for bait-prey interactions. To capture additional evidence of interaction between prey proteins, we reanalyzed AP-MS, RNA pulldown, and Proximity Labeling datasets using our weighted matrix model approach (WMM)^17,23^ to identify pairs of proteins seen in these large datasets more often than random (see methods). This feature has been extremely valuable at identifying novel interactions in previous work^16,17^.

### Construction of machine learning model to identify protein interactions

We next build a machine learning model using evidence features to identify pairs of proteins that interact in a complex. Once we constructed our feature matrix, we label the pairs of proteins with our training set derived from manually curated protein complexes in the Complex Portal^21^. To generate a training and test set, we randomly split protein complexes from Complex Portal and remove any pairs that overlap. We next label our feature matrix as either being previously seen in a training complex together (i.e. positive label), seen in different training complexes (i.e. negative label), or not seen at all (i.e. unknown) (see methods). We then use the feature matrix labeled with training data to build a ML classifier. For this step we use AutoGluon^24^, which automates model and hyper-parameter selection across 13 machine learning models. We empirically evaluated several evaluation metrics including accuracy, precision, and F1 score, and identified accuracy as having the optimal performance (Fig S1A). The AutoGluon step builds a weighted ensemble model based on the top performing classifiers. We then generate a network consisting of ∼26 million protein pairs where all pairs of proteins are classified using the final weighted ensemble model. The model produces a confidence score for each pair of proteins representing the estimated probability of the pair interacting based on the evidence.

We then evaluate our final model using a precision recall analysis evaluated on a leave out test set of protein interactions (Fig 2A). We compare our current hu.MAP3.0 protein interaction network against our prior protein interaction network, hu.MAP2.0, as well as the Bioplex3 interaction network. We see our current hu.MAP3.0 outperforms the other networks, increasing in both precision and recall, with an area under the precision recall curve (AUPRC) of 0.68.

**Figure 2.**
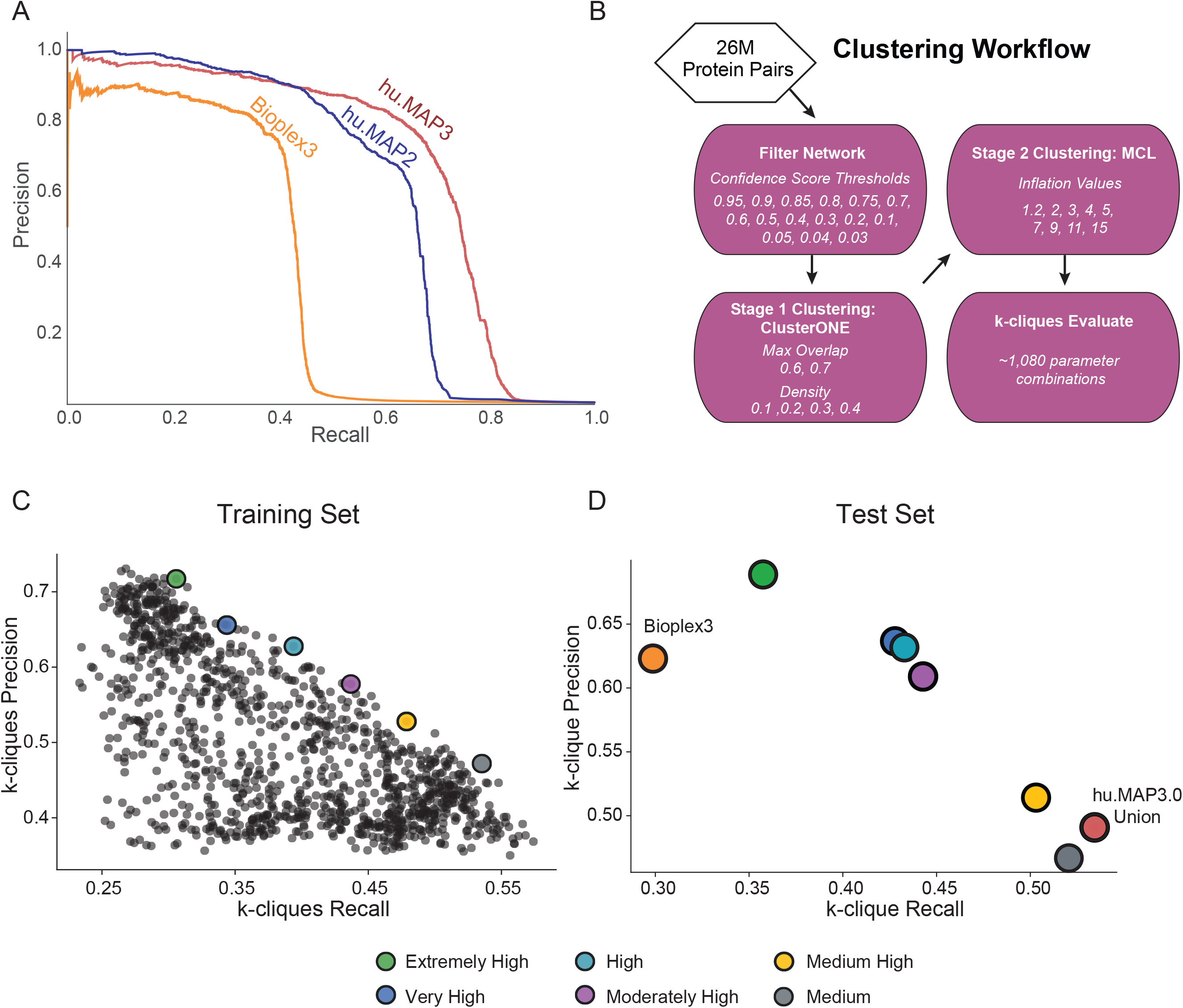
hu.MAP3.0 outperforms previous complex maps. (**A**) Precision recall (PR) analysis on a test leave out set of gold standard co-complex interactions demonstrates hu.MAP3.0 is more accurate and comprehensive than alternative models, hu.MAP2 and Bioplex3. (**B**) Clustering workflow to identify optimal clustering parameters. (**C**) Scatterplot of *k*-clique precision recall measures for >1,000 clustering parameters. *k*-clique evaluation in (**C**) is done on a training set of gold standard complexes to identify parameters that produce confident protein complexes. We observe a trade off between precision and recall. We selected six parameter sets that represented “extremely high” confidence (green) to “medium” confidence (gray) complexes. (**D**) *k*-clique evaluation of selected sets of complexes using a test leave out set of gold standard complexes. Consistent with (**C**), we observe the same tradeoff between precision and recall, in addition to maintaining ordering of sets of complexes. Additionally, the union of all hu.MAP3.0 complexes from the various confidence levels improves on recall. Extremely High, Very High and High confident complexes improve in both precision and recall over Bioplex3 complexes.

Bioplex3 and hu.MAP2.0 have an AUPRC of 0.38 and 0.61, respectively. To assess how well the confidence score generated by our model reflects the true precision of the network, we compare the confidence score to precision as determined by the leave out test set. In Figure S1B-C, we see the confidence score closely reflects the true precision. We also see the relationship between confidence score and test set label (true or false interaction) in Figure S1D. Here we see the majority of true interactions have substantially higher confidence scores and the vast majority of false interactions have very low confidence scores.

### Identification of protein complexes within protein interaction network

We next cluster the full protein interaction network to identify protein complexes. This is done using a two stage clustering approach (Fig 2B). We first use the ClusterOne algorithm^25^ followed by the MCL algorithm^26^ which both have free parameters to define. To determine optimal clusters, we test combinations of parameters in both algorithms as well as determine a confidence threshold for the input protein interaction network. We use each parameter combination to generate a set of clusters which are then evaluated against a training set of known protein complexes. Figure 2C shows the evaluation of all sets of clusters from 1,080 parameter combinations on the set of curated training complexes. Each set of clusters is evaluated using the *k*-clique precision and recall measures which globally compares a set of clusters against a benchmark set^17^. In this evaluation (Fig 2C), we see a tradeoff between precision and recall, where some cluster sets have high precision low recall, other cluster sets have lower precision high recall, and others are not optimized for either. We select six clustering sets along this boundary to combine into our final set of hu.MAP3.0 predicted complexes. We rank each set of clusters according to their precision value, considering the highest precision as “Extremely High” confident, followed by “Very High”, “High”, “Moderately High”, “Medium High”, and “Medium”. The last of which, “Medium” has the highest recall of all other cluster sets. The full set of hu.MAP3.0 complexes can be found in Table S1. We then evaluate all six cluster sets and their union on a benchmark of curated leave out test complexes (Fig 2D). We see that all six cluster sets have a consistent order in terms of their tradeoff between precision and recall, with “Extremely High” clusters having the highest precision and lowest recall, and the “Medium” clusters having the highest recall and lowest precision. Specific complexes were seen multiple times throughout the confidence values. We therefore assigned each final complex the highest confidence level in which the complex was seen. Figure S2A shows the distribution of complex size relative to complex confidence levels. We see complex size increase with lower levels of confidence (i.e. higher recall). This is likely due to core members of complexes being confidently identified in high confidence levels and additional auxiliary subunits added to complexes in lower confidence clusters. Total number of complexes is highest in our highest confidence level (Fig 1D). Figure 1B shows our highest confidence level has broad coverage of the proteome.

We observe the total number of protein-protein interactions (PPIs) per confidence level rise as we decrease in confidence, again suggesting that high confidence complexes are smaller representing core subunits (Fig 1C).

We also evaluate the union of all cluster sets, and observe it has the highest recall over all the individual cluster sets and see hu.MAP3.0 complexes outperform Bioplex3 clusters (Fig 2D). We also compare the coverage of proteins, protein interactions, and complexes and see hu.MAP3.0 increases comprehensiveness over both Bioplex3 and hu.MAP2.0 (Fig 1B-D). Finally, we observe nearly 40% of complexes are enriched with an annotation including from Gene Ontology (GO)^27^, KEGG^28^, Reactome^29^, CORUM^30^, and Human Phenotype Ontology (HP)^31^ (Fig S2B, Table S2).

### Protein covariation provides orthogonal evidence for hu.MAP3.0 complexes

Protein covariation analysis, also known as coexpression or co-regulation analysis, identifies proteins exhibiting coordinated changes in abundance across biological conditions. We and others have previously shown that protein covariation analysis is a powerful approach to capture functional relationships between proteins and provide novel insights into proteome organization^19,20,32–36^. For our protein covariation resource, ProteomeHD, we re-processed thousands of mass spectrometry runs that had been manually selected to reflect protein abundance changes in response to various biological perturbations^19^. For example, typical experimental designs included comparisons of mutant to wildtype cells, and drug *vs* control treatments. We then used a decision-tree - based algorithm to determine likely functional relationships between proteins based on correlated expression patterns. Therefore, hu.MAP and ProteomeHD are conceptually similar. Both approaches integrate thousands of proteomics experiments using machine-learning to infer protein - protein associations in a data-driven manner. However, they are also highly complementary as they rely on fundamentally different types of evidence and raw data. While hu.MAP integrates experiments that determine physical interactions, protein covariation analysis draws on expression data from an entirely distinct set of mass spectrometry experiments, highlighting functional relationships based on covariation rather than physical interaction.

Despite these differences, we and others have previously found that many strongly covarying proteins are in fact subunits of the same protein complex^19,20,34,37^, suggesting that many complexes have strong covariation signatures. Therefore, we reasoned that protein covariation could provide orthogonal evidence to validate, enrich and annotate our human protein complex map. To test this we combined protein-protein interaction probabilities from hu.MAP3.0 with covariation scores from ProteomeHD.2 (ProHD.2), a recently developed successor of ProteomeHD (Fig 3A). In comparison to the first iteration of ProteomeHD, ProHD.2 incorporates a larger number of biological conditions and uses an improved, supervised machine-learning strategy to determine proteins whose abundance covaries across these conditions^38^ (see methods). In total, 15 million protein pairs are covered by both hu.MAP3.0 and ProteomeHD.2 (Fig S3A). To assess if the datasets are indeed complementary, we compared them to high-quality, experimentally determined pairwise interactions from the OpenCell project^6^, which neither hu.MAP3.0 nor ProHD.2 used as input. Indeed, protein pairs with high association scores in both hu.Map3.0 and ProHD.2 are much more likely to be captured by OpenCell than random pairs (29% vs 0.3%, respectively), and significantly more likely than pairs captured by only one of the two approaches (Fig S3B). This suggests that protein covariation provides orthogonal evidence to support protein interactions in hu.MAP3.0.

**Figure 3.**
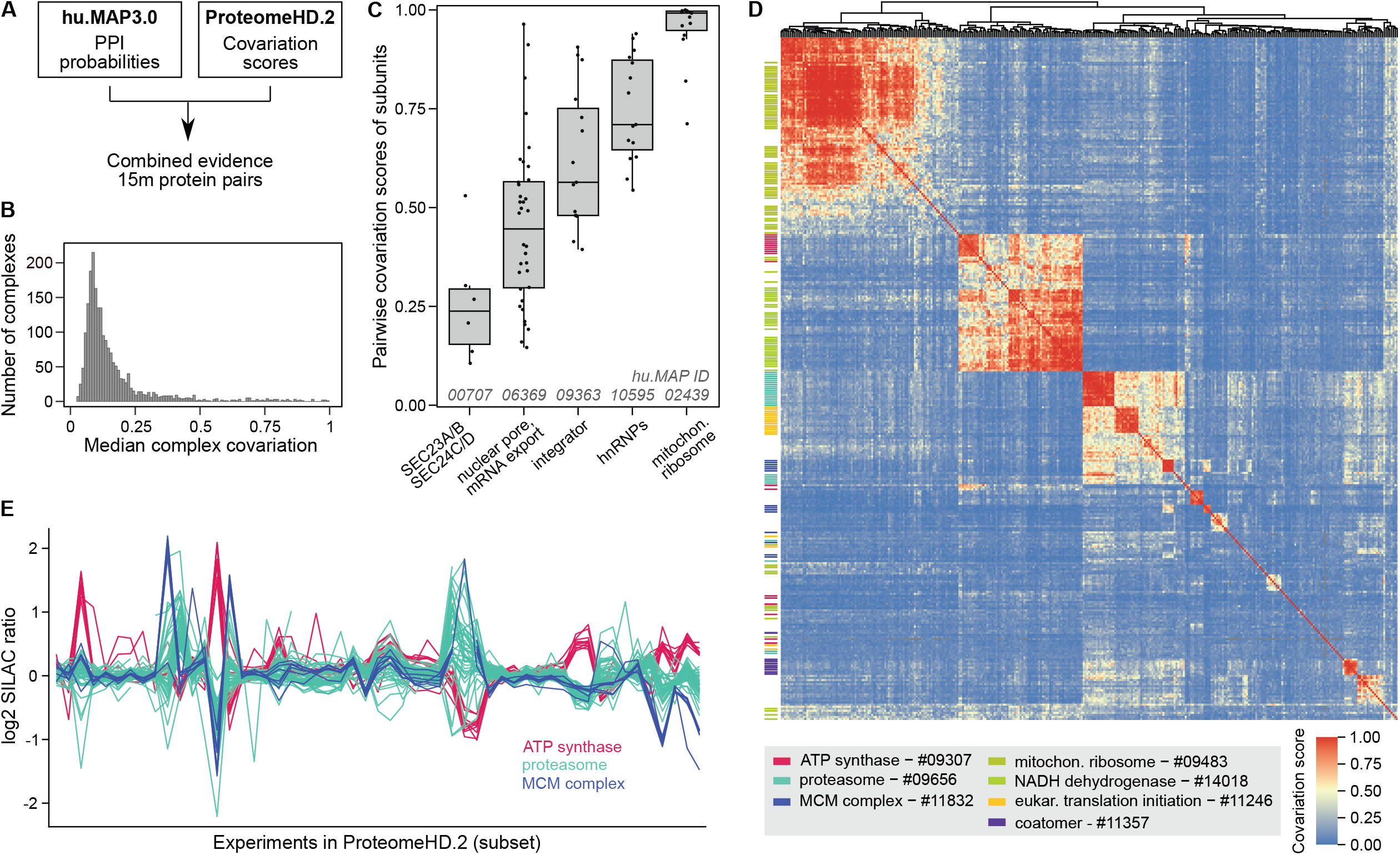
Integrating hu.MAP3.0 with coexpression data reveals complementary insights into protein complex biology. (**A**) Workflow of pairwise protein-protein interaction probabilities (hu.MAP3.0) with pairwise protein covariation probabilities (ProteomeHD.2). Protein identifiers were combined at the protein-level, with only common pairs between datasets used for downstream analysis. (**B**) Histogram showing median covariation of subunits for hu.MAP3.0 complexes. Complexes with fewer than 4 subunits, a hu.MAP3.0 confidence score below 4, or less than 50% coverage in ProteomeHD.2 were excluded. (**C**) Boxplots of pairwise subunit covariation scores for five example complexes, selected to represent diverse degrees of complex-level covariation. (**D**) Heatmap showing hierarchical clustering of protein covariation scores for proteins assigned to hu.MAP3.0 complexes. Subunits of selected complexes are highlighted as a rug plot on the left, indicating that many complex subunits cluster together. Proteins are plotted in the same order on both axes, revealing covariation between different complexes. Only complexes with 50% coverage in ProteomeHD.2 and median covariation score of 0.45 were included. (**E**) Log2 fold-changes (SILAC ratios) across a subset of perturbation experiments from ProteomeHD.2 for subunits belonging to three selected complexes.

### Covariation provides complementary insights into complex biology

We next asked what the combined hu.MAP3.0 and ProHD.2 data could tell us about the nature of protein complex expression. We find that complexes show varying degrees of subunit covariation (Fig 3B). Some hu.MAP3.0 protein complexes are tightly co-expressed modules with all subunits showing highly coordinated abundance changes, such as the mitochondrial ribosome and a nuclear ribonucleoprotein complex (Fig 3C). Other complexes, such as the nuclear pore and integrator complexes, have weaker but still very clear coexpression signatures. However, many complexes show low subunit covariation. There could be technical explanations for this, such as a failure of ProHD.2 to capture covariation patterns for some proteins (false negatives in ProHD.2) as well as potential complex prediction errors (false positives in hu.MAP3.0). However, we find that the degree of complex covariation correlates only mildly with hu.MAP3.0 confidence levels (Fig S4), suggesting that poor subunit covariation is not simply the result of potential complex prediction errors. Low covariation scores may also be observed for biological reasons, for example if complex subunits have additional, non-overlapping functions outside of the complex. In the case of a hu.MAP3.0 complex consisting of SEC23A, SEC23B, SEC24C and SEC24D (Fig 3C), a low subunit covariation could reflect the fact at SEC23A / SEC23B and SEC24C / SEC24D are paralogues, respectively^39^, which could potentially form mutually exclusive Sec23-24 dimers. We explore the possibility to use our data to systematically detect mutually exclusive subunits below. Taken together, these results suggest that, while a high covariation score supports physical interaction evidence, a low covariation score is more difficult to interpret and does not necessarily imply a lack of physical interaction.

### Protein covariation captures associations between and within complexes

Unlike traditional protein-protein interaction mapping, protein covariation can identify functionally associated proteins even when they do not physically interact. This is because proteins involved in related biological processes may exhibit coordinated abundance changes despite the lack of direct physical contact. For example, we have shown that the peroxisomal protein PEX11β covaries with proteins involved in mitochondrial respiration^19^. Now, we leverage this property to assess inter-complex covariation, determining which complexes display similar expression patterns across the ProHD.2 dataset. This analysis enabled us to construct a map connecting dozens of hu.MAP3.0 complexes (Fig 3D). It identifies associations between functionally related complexes, such as the mitochondrial respiratory complexes NADH dehydrogenase (complex I) and ATP synthase (complex V). However, these two complexes do not co-vary with the mitochondrial ribosome, indicating that the detection of inter-complex associations is function-specific and does not merely capture subcellular localisation. We observe strong covariation between two hu.MAP3.0 complexes enriched in translation initiation factors and proteasomal proteins, respectively. This association may reflect a role of the proteasome in translation quality control^40^. However, some inter-complex associations are more difficult to interpret. For example, the proteasome shares a broadly similar expression profile with subunits of the MCM replication factor complex (Fig 3D, E).

We next explored if covariation analysis could reveal substructures within individual protein complexes, using the proteasome as a well-annotated example. The 26S proteasome is a large 30+ subunit protein complex responsible for protein degradation and maintaining protein homeostasis. The full complex is made up of defined subcomplexes including a 20S core and 19S regulatory particles^41^. The hu.MAP3.0 complex huMAP3_09656.1 includes these core and regulatory subunits of the proteasome, along with tissue-specific core subunits and several assembly chaperones, activator complexes and other proteins known to associate with the proteasome. The degree of covariation between members of this complex reflects known biological properties (Fig 4A). Subunits from the 20S core and the 19S regulatory particle exhibit strong covariation within their respective subcomplexes, with slightly lower covariation observed between these two substructures. The immunoproteasome-specific subunits PSMB8-10, which replace PSMB5-7 in immune cells, show weaker covariation with other core subunits compared to their canonical counterparts^41^. The two subunits of the PA28αβ activator (PSME1, PSME2) strongly covary with each other and with multiple proteasome subunits, but demonstrate weak covariation with the PA28γ activator (PSME3), which, despite structural similarity, has a distinct biological role^41^. Additionally, the proteasome assembly chaperone PAC3 (PSMG3), which forms a heterodimer with PAC4 (PSMG4), interacts preferentially with the α5 core subunit (PSMA5), as reflected in the higher covariation scores with these proteins^42^. Furthermore, covariation data support established proteasome interactions with other proteins, such as AKIRIN2, which plays a role in nuclear import of proteasomes^43^, and UCHL5, a proteasome-associated deubiquitinase^44^.

**Figure 4.**
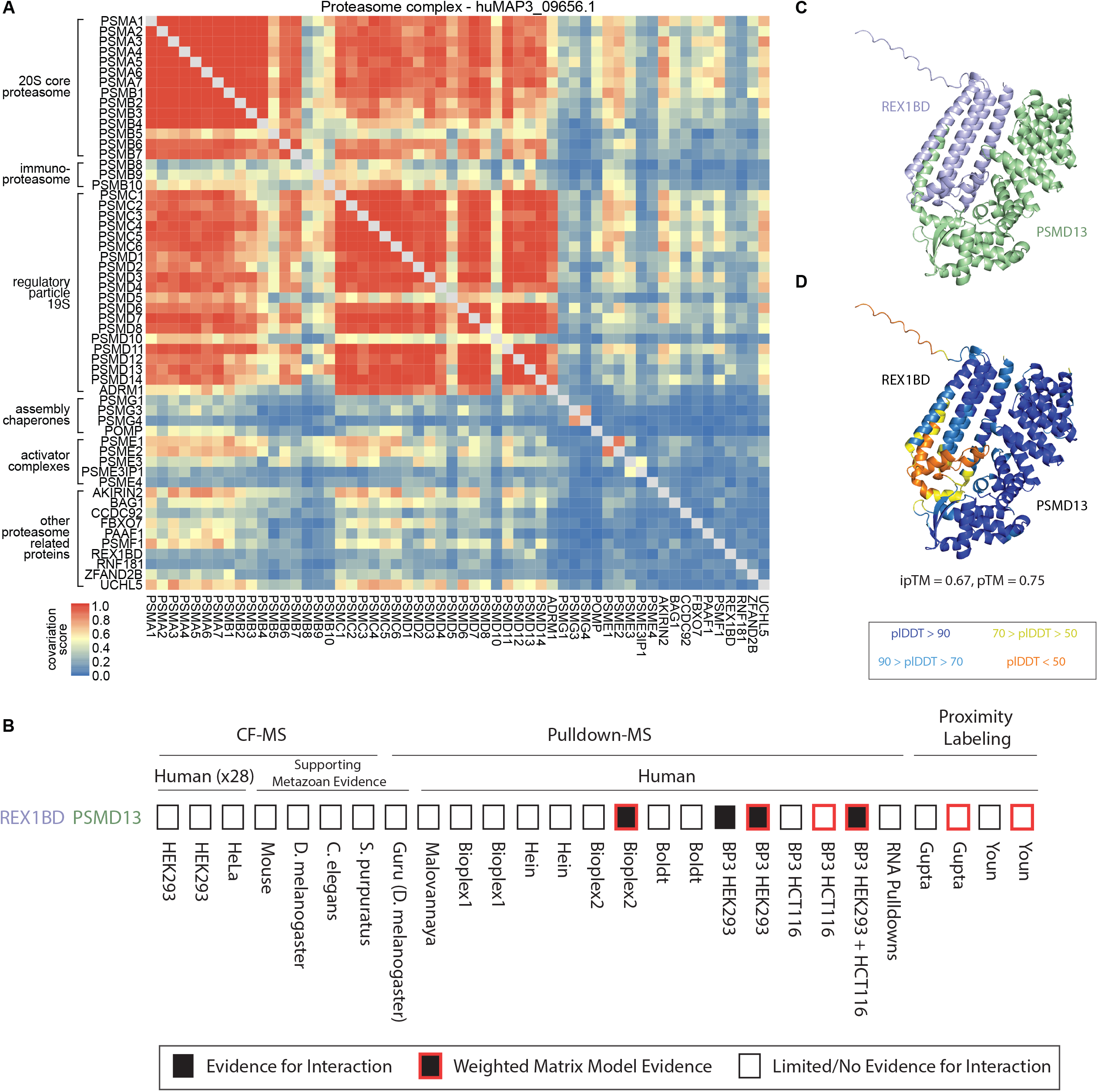
Covariation identifies modules of proteasome and novel interactor of 19S regulatory lid. (**A**) Proteasome heatmap shows pairwise covariation score. (**B**) Graphical representation of evidence for protein interaction. (**C**) AlphaFold3 model of uncharacterized protein REX1BD (blue) and PSMD13 (green) subunit of proteasome lid. (**D**) Same model in (**C**), colored by pLDDT confidence score. Color scale for pLDDT is from blue (very high confidence) to orange (very low confidence).

### Uncharacterized protein REX1BD interacts with proteasome lid subunits

Within the proteasome hu.MAP3.0 complex, we identify potential members not previously known to associate with the full complex. Specifically, we identified REX1BD, an uncharacterized protein as a high confidence co-complex interactor of the 19S proteasome lid, including 19S lid subunits PSMD13 (hu.MAP3.0 score = 0.99), PSMD14 (0.99), and PSMD7 (0.99). We also find evidence for physical association between PSMD14 and REX1BD through immunoprecipitation studies^45^. Figure 4B shows the evidence from individual proteomic experiments and analysis for REX1BD-PSMD13 interaction which included both pulldowns and weighted matrix models of the pulldown compendium. Additionally, we find REX1BD and PSMD13 demonstrate coordinated abundance changes in ProteomeHD.2 (Fig 4A). Their covariation score of 0.43 places the pair in the top 0.6% of all associations scored by ProHD.2. To provide further evidence for this interaction, we used AlphaFold3 to model the three dimensional structure of REX1BD and PSMD13. AlphaFold3 produced a confident model of the two proteins (pTM=0.75) including a confident interaction interface (ipTM=0.67) (Fig 4C,D).

These results suggest a potential role of REX1BD in the proteasome degradation pathway.

### hu.MAP3.0 provides testable hypotheses to understudied proteins

Building upon our identification of the uncharacterized REX1BD as interacting with the 19S proteasome, we next searched for functional hypotheses for other uncharacterized proteins. Thousands of human proteins, many from essential genes and of biomedical importance, remain understudied^46^. We and others have used protein complex maps as an unbiased way of transferring function annotations to uncharacterized proteins as well as identifying novel disease genes^16,17,47–49^.

As an example of identifying novel disease genes, we find a novel interaction between disease gene IER3IP1 and an uncharacterized protein TMEM167A (huMAP3_00596.1, hu.MAP3.0 score = 0.992) (Fig 5A,B). IER3IP1 is associated with brain development and, when mutated, gives rise to developmental abnormalities including microcephaly, epilepsy, and neonatal diabetes syndrome (MEDS)^50–52^. At the cellular and molecular level, IER3IP1 localizes to the Golgi, regulates the endoplasmic reticulum (ER) and is involved in secretion of extracellular matrix proteins^53^. We see evidence for this interaction from pulldown experiments as well as weighted matrix model analysis of pulldowns (Fig 5C). Both IER3IP1 and TMEM167A are predicted to be transmembrane^54^. Like IER3IP1, TMEM167A also localizes to the Golgi and is involved in protein secretion^55^. The ProteomeHD.2 network identifies a strong covariation between these two proteins (0.78) suggesting they are regulated as a module. Using AlphaFold3^56^, we predicted a confident 3D model of IER3IP1 interacting with TMEM167A (ipTM = 0.77, pTM = 0.8) (Fig 5A,B). Finally, we modeled two known missense mutations in IER3IP1 causative for MEDS in human patients (Val21Gly and Leu78Pro)^52^. Figure 5A shows these two mutations are at the interface of IER3IP1’s interaction with TMEM167A suggesting a molecular mechanism for this disease.

**Figure 5.**
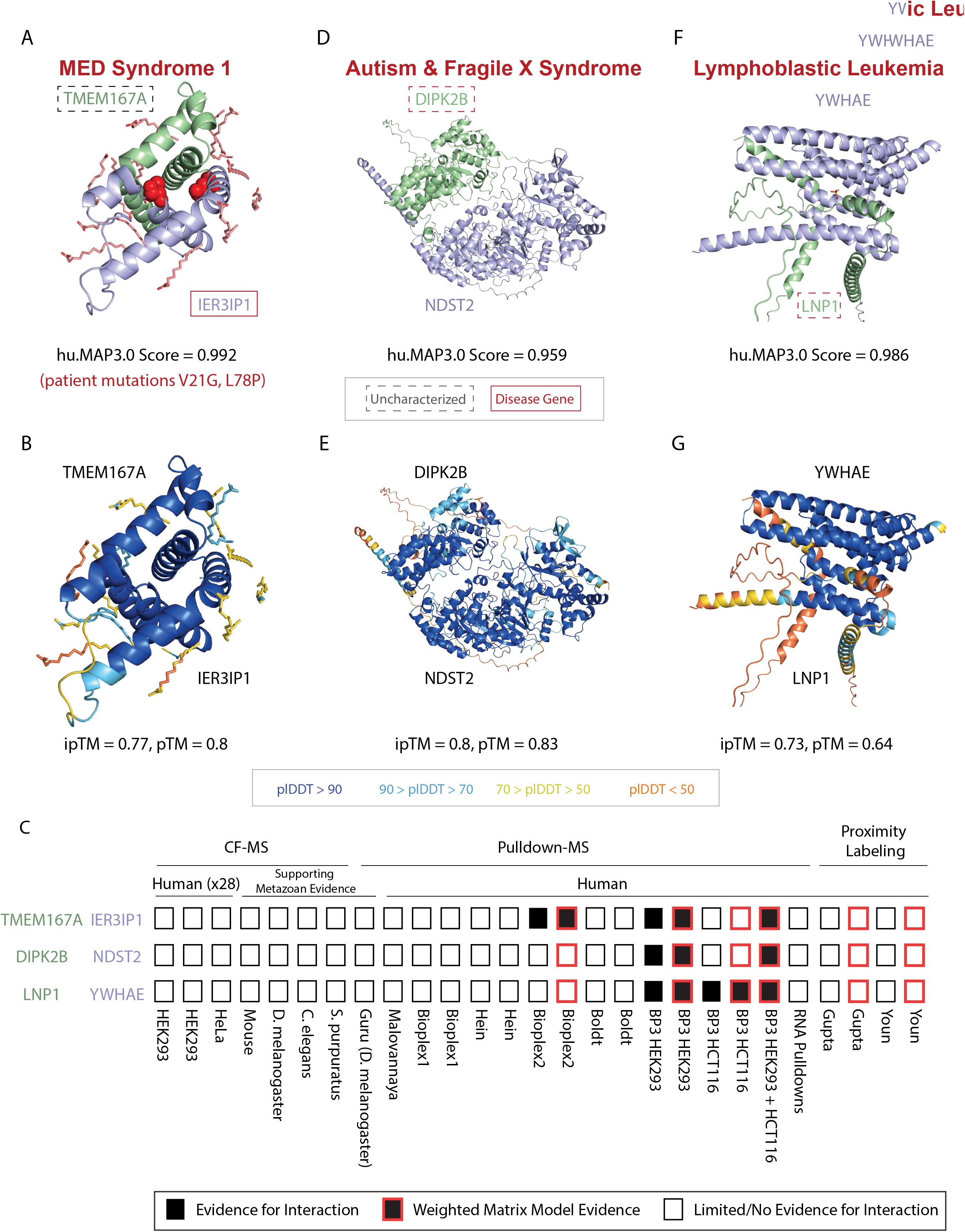
**Physical associations with uncharacterized proteins.** (**A**) AlphaFold3 model of uncharacterized protein TMEM167A (green) and IER3IP1 (blue), a protein associated with MED Syndrome. Known patient mutations are shown in red spheres at the interaction interface. (**B**) Same model in (**A**), colored by pLDDT confidence score. (**C**) Graphical representation of evidence for protein interactions. (**D**) AlphaFold3 structural model of autism and Fragile X syndrome protein, DIPK2B (green), and NDST2 (blue) colored by chain. (**E**) Same model in (**D**), colored by pLDDT confidence score. (**F**) AlphaFold3 structural model of lymphoblastic leukemia protein, LNP1 (green), and YWHAE (blue). (**G**) Same model in (**F**), colored by pLDDT confidence score. Uncharacterized genes are highlighted with a dashed box. Disease genes are highlighted with a red box. Color scale for pLDDT is from blue (very high confidence) to orange (very low confidence).

Given this example, we next searched annotated hu.MAP3.0 complexes for the least characterized proteins to identify potential functions based on the “guilt-by-association” principle. Specifically, we searched for uncharacterized proteins (UniProt Annotation Score <= 3) which were physically associated with complexes statistically enriched for function annotations (Figure S2B). Our analysis identifies potential functions for 472 of the least characterized human proteins (see Table S3). Several of these functionally uncharacterized proteins are associated with human diseases including ciliary dyskinesia (CLXN^57^), Lui-Jee-Baron syndrome (SPIN4^58^), lymphoblastic leukemia (LNP1^59^, SLX4IP^60^), lethal skeletal dysplasia (TMEM263^61^), and association to autism and fragile X syndrome (DIPK2B^62^).

The autism-associated protein, DIPK2B, is a member of the FAM69 family of protein kinases and has a signal peptide which localizes the protein to the extracellular region^63^. We identified DIPK2B as being co-complex with two proteins NDST2 and NDST3 (huMAP3_07022.1) involved in glycosaminoglycan biosynthesis (KEGG:00534). NDST2 and NDST3 are bifunctional enzymes which catalyze N-deacetylation and N-sulfation of glucosamine in the synthesis of heparin sulfate of the extracellular matrix^64,65^. Heparin sulfate deficiencies are thought to be causative for autism disease^66^. To provide further evidence for this interaction, we modeled the interaction between DIPK2B and NDST2 (hu.MAP3.0 score = 0.959) using AlphaFold3 in which we obtained a confident structural model of the heterodimer (ipTM = 0.8) (Fig 5D,E). Taken together, we propose that DIPK2B plays a role in maintaining heparin sulfate of the extracellular matrix and this is linked to its association with autism disorder.

We also identify LNP1, an uncharacterized protein, as associated with members of the 14-3-3 complex (huMAP3_06971.1). LNP1 is known to fuse with NUP98 in genome rearrangements of leukemia patients while the 14-3-3 complex is involved with many signaling pathways and associated with cancer^67^. LNP1 has a known phosphoserine site at Ser114^68^ in a motif reminiscent of 14-3-3 binding (KFpSESF vs RXY/FXpSXP^69^). Provided this, we used AlphaFold3 to model the interaction between LNP1 and YWHAE, a 14-3-3 subunit which had the highest hu.MAP3.0 score 0.986 to LNP1. Figure 5F-G shows LNP1 and YWHAE confidently interact with a phosphorylated Ser114 interacting at the interface. This provides further evidence of LNP1’s association with the 14-3-3 complex.

In total, our guilt-by-association transfer of function annotations results in a >70% increase from hu.MAP2.0^16^ (472 vs 274) in terms of the total number of uncharacterized proteins annotated.

We find our analysis provides strong testable hypotheses for these understudied proteins and should guide followup studies.

### Identification of mutually exclusive subunits in hu.MAP3.0 complexes using AlphaFold structural models

Protein complexes often have modular components which, depending on the subunit present, alter the complex’s function. When these multiple subunits exist, the subunits potentially compete for a binding interface and therefore cannot exist together in the same complex at the same time (i.e. mutually exclusive). Well known examples include replacement of PSMB5 for PSMB8 in the immunoproteasome^41^ (see above), ARID1A and ARID2 stabilizing the core of the chromatin remodeling PBAF complex, altering gene targeting^70^, and SINA and SINB in the Sin3 complex^71^.

Three dimensional structures can be used to identify mutually exclusive subunits by observing overlapping interfaces with a third common subunit. We first examine whether we can use information from experimental structures to determine mutually exclusive protein subunits.

COPS7A and COPS7B are well characterized mutually exclusive subunits as determined by genetic and biochemical data and differentially alter the function of the COP9 Signalosome, an essential regulator of the ubiquitin degradation pathway^72–74^. Our hu.MAP3.0 pipeline identifies a COP9 Signalosome complex (humap3_05041.1) (Fig S5A) with strong interactions between many of the subunits including COPS7A and COPS7B (hu.MAP3.0 score = 0.993). The structures of COP9 Signalosome with each COPS7A and COPS7B, separately, have each been experimentally determined^75,76^. Upon alignment and inspection of each structure, we identify COPS7A has the same binding interface with the rest of the core COP9 Signalosome complex as COPS7B (Fig S5B). This strongly confirms the subunits are mutually exclusive. Given this, we next turn to the ProteomeHD.2 network to ask if co-expression can discriminate between variants of the complex. Figure S5C,D shows the ProteomeHD.2 co-expression network of the COP9 Signalosome. We observe strong co-expression among core subunits, low co-expression among related proteins (NEDD8, FBXO17, APPBP2, KLHDC3) to core subunits, and moderate co-expression of COPS7A and COPS7B to core subunits. Alternatively, we see weak co-expression between COPS7A and COPS7B (ProteomeHD.2 score = 0.25) suggesting subunit expression patterns have altered to avoid conflicting functions and may be an indicator of mutual exclusion.

We next ask if high accuracy structural models (e.g. AlphaFold) are capable of identifying mutually exclusive subunits similar to experimental structures. Figure S5E shows an example of a hu.MAP3.0 identified multifunctional protein complex, BRISC/BRCA1-A complex (huMAP3_08508.1), which exhibits mutually exclusive proteins ABRAXAS1 and ABRAXAS2^77^. From experimental structures^78^ and AlphaFold3 models (Figure S5F), we can see that ABRAXAS1 and ABRAXAS2 subunits are mutually exclusive as they overlap in their three dimensional coordinates and occupy the same interface with BRCC3. We observe that ABRAXAS1 and ABRAXAS2 have limited co-expression (ProteomeHD.2 score = 0.096), consistent with the idea that the two proteins avoid competing for the same binding interface using different protein expression patterns. ABRAXAS2 is therefore identified as a member of the BRISC complex (Figure S5G) and ABRAXAS1 is a member of the BRCA1-A complex (Figure S5H). Additionally, we observe an uncharacterized protein, C9orf85, as being physically associated with core subunits of the BRISC/BRCA1-A complex. We see strong evidence for a physical interaction between C9orf85 and ABRAXAS2 (hu.MAP3.0 score = 0.989) but limited evidence for C9orf85 being physically associated with ABRAXAS1 (hu.MAP3.0 score = 0.0).

This points to C9orf85 as a member of the BRISC complex and potentially contributes to its specific function (Figure S5G). As shown in these examples, it is highly beneficial to have three dimensional structures of protein complexes to evaluate mutually exclusive subunits. Several studies have applied AlphaFold2 / AlphaFold-multimer workflows to determine structures of thousands of protein interactions^18,79^. We next utilize these compendiums of pairwise structural models to systematically identify potential mutually exclusive pairs in hu.MAP3.0 complexes (Fig 6, center panel). We first filtered the structural models to determine confident predictions. We then identified confident models that share a common subunit and aligned on the common subunit. Finally, we evaluated the interfaces of proteins with the common subunit for potential overlap. If an overlap of proteins exists, the proteins are labeled mutually exclusive. Alternatively, if no overlap exists at the interface the interaction is labeled structurally consistent (see methods).

**Figure 6.**
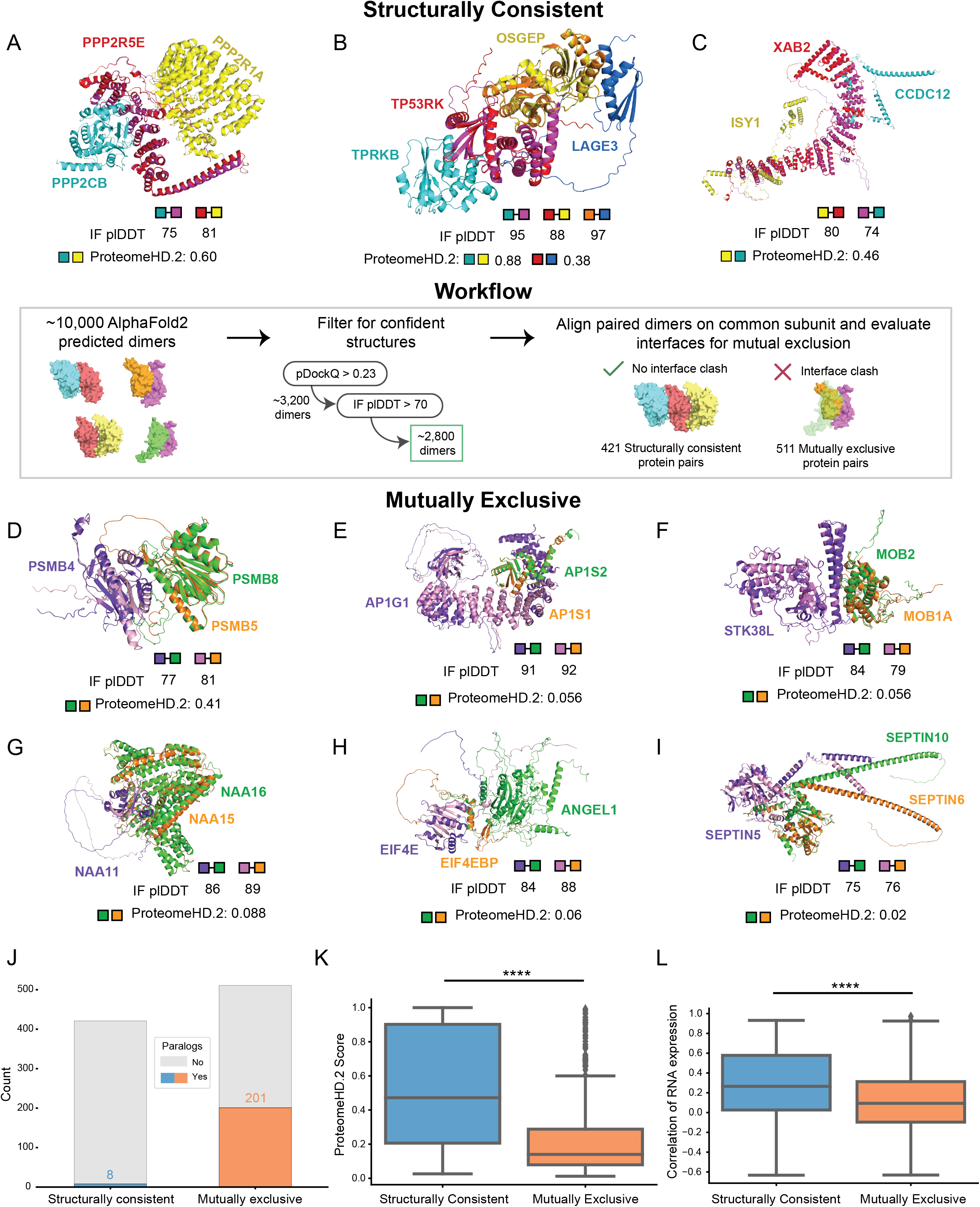
Systematic structural analysis reveals mutually exclusive interactions. (**Center panel**) Overview of the workflow utilizing AlphaFold2 models to identify structurally consistent and mutually exclusive interactions within hu.MAP 3.0 protein complexes. **Structurally Consistent** (**A, B,** and **C**), Examples of interactions are shown with aligned AlphaFold2 models, where the common subunit (magenta and red) and the unique proteins (cyan and yellow) form structurally consistent interactions. The interfaces between the unique protein and common proteins do not overlap. IF_pLDDT is displayed for each dimeric interface to indicate interface quality. The ProteomeHD.2 is displayed for the unique proteins to indicate their covariation across tissues. The Keops complex (**B**) has two structurally consistent trimers TPRKB-TP53RK-OSGEP and TP53RK-OSGEP-LAGE3, which is displayed as a tetramer. **Mutually Exclusive** (**D - I**), In contrast, mutually exclusive interactions are shown with aligned AlphaFold2 models, where the common protein subunit (light pink and purple) interacts with the unique proteins (green and orange) and the interfaces overlap. As before, interface quality metrics and the ProteomeHD.2 score for the unique protein subunits are shown. (**J**) Stacked barplot showing the prevalence of paralogs present in structurally consistent and mutually exclusive interactactions, with a higher frequency of paralogs found in mutually exclusive interactions. (**K**) Boxplot of the global distribution of ProteomeHD.2 scores between structurally consistent and mutually exclusive interactions, with structurally consistent interactions exhibiting stronger covariation compared to mutually exclusive interactions (Welch’s *T*-test for independent samples, \*\*\*\**P* < 0.0001; box corresponds to quartiles of the distribution, line in box corresponds to median, whiskers extend to the first data point inside the first/third quartile plus 1.5 × interquartile range (IQR), respectively). (**L**) Boxplot comparing the correlation of RNA expression across different tissues for the unique proteins in structurally consistent versus mutually exclusive interactions (two-sided Mann-Whitney *U*-Test, \*\*\*\**P <* 0.0001; boxplot descriptions same as **K**).

Using this method we identify 511 mutually exclusive and 421 structurally consistent pairs of proteins in hu.MAP3.0 complexes (Table S4).

Figure 6A-C shows examples of structurally consistent interactions identified by our workflow. This highlights unique interfaces for pairs of proteins sharing a common third protein. These models include the PP2A complex with a structural model for the catalytic subunit PPP2CB and interactions from the KEOPS complex, which has been previously modeled in yeast^80^ and human using homology modeling^81^. Further, we identify members of the spliceosomal complex with an uncharacterized protein, CCDC12. We observe high pairwise ProteomeHD.2 scores of structurally consistent interactions suggesting they are co-expressed as protein modules.

Figure 6D-I shows examples of proteins we labeled as mutually exclusive interactions. We identify positive control examples PSMB5 and PSMB8 known to substitute for each other in the proteasome^41^ (Fig 6D). We also identify paralog subunits of the heterotetrameric adaptor protein I complex (AP-1), AP1S1 and AP1S2, to be mutually exclusive (Fig 6E). This finding is supported by AP1S1 and AP1S2 having distinct, non-redundant functions that lead to different disease pathologies when mutated^82,83^ as well as differential RNA expression patterns across tissues (Pearson’s correlation coefficient = -0.08, Table S4)^84^.

We also identify key interactions that regulate protein function and cell signaling. For example, STK38L’s kinase activity has been shown to be regulated by both MOB1A and MOB2, where MOB1A stimulates STK38L phosphorylation activity and MOB2 inhibits it^85^. Further, biochemical data shows physical interactions between STK38L and both MOB1A and MOB2^86^. A crystal structure also exists for STK38L and MOB1A^87^. It has been previously shown that MOB1A and MOB2 compete for binding on STK38^88^, a paralog of STK38L. Figure 6F displays structural models showing MOB1A and MOB2 interact at the same interface on STK38L providing structural support to their mutual exclusivity.

Additionally, we characterized interactions of the Nat A complex, which is responsible for post-translation N-terminal acetylation^89^. NAA11 is an understudied paralog of the well characterized Nat A subunit NAA10^90^. NAA16 is thought to substitute for NAA15 and carry out N-terminal acetylation in NAA15-knockdown experiments^91^. We identify NAA15 and NAA16 as having overlapping interfaces with NAA11 (Fig 6G) supporting the idea that NAA16 can substitute for NAA15.

Furthermore, we observe interactions within the eIF4F translation initiation complex, which regulates cap-dependent mRNA translation^92^. EIF4E is subject to translational repression, through interactions with EIF4BP1, which prevent its incorporation to the eIF4F complex^93^. Biochemical data shows ANGEL1 as an interaction partner of EIF4E, and uses a similar binding motif as EIF4EBP1^94^. Our structural modeling reveals overlapping interfaces between EIF4BP1 and ANGEL1 when bound to EIF4E, supporting the mutual exclusivity of these interactions (Fig 6H). Despite their mutual exclusivity, overexpression of either protein does not reduce binding of the other. This is likely due to their distinct cellular localizations where EIF4EBP1 localizes to the cytoplasm and ANGEL1 is confined to the golgi apparatus and endoplasmic reticulum^94^. This suggests regulation of subcellular localization plays a key role in maintaining this exclusivity.

We also identify interactions of septin complexes, which form various filament-forming, hetero-oligomeric complexes and are crucial components of the cytoskeleton, such as protein scaffolding and diffusion barriers^95^. The SEPTIN2-SEPTIN6-SEPTIN7 complex is the most studied and abundant septin assembly^96^. However, based on sequence homology, previous research suggests that subunits within this complex are often substituted^97^. For example, SEPTIN5, SEPTIN4, or SEPTIN1 can replace SEPTIN2, while SEPTIN8, SEPTIN10, or SEPTIN11 can replace SEPTIN6. Pull down experiments further suggest interactions of SEPTIN5 with SEPTIN6 and SEPTIN10^3^, though many other septins are detected as well, likely reflecting their filamentous assembly. Our findings uniquely reveal mutual exclusivity between SEPTIN 10 and SEPTIN6 when binding to SEPTIN5 suggesting they compete for a shared interface (Fig 6I).

We next asked if proteins involved in mutually exclusive interactions are enriched with Gene Ontology annotations. This may point to certain biological systems whose functions are more readily regulated by substitutions of subunits. To determine potential functional enrichment signatures of mutual exclusivity, we analyzed Gene Ontology annotations of all proteins identified in our mutually exclusive dataset (Fig S6A). We found enrichment in “vesicle-mediated transport”, “cytoskeletal organization”, and “kinase activity”. These align with our highlighted examples above, AP1 complex (vesicle transport), septin interactions (cytoskeleton) and STK38L-MOB1A-MOB2 interactions (kinase activity). Overall this suggests these biological systems reuse core components and alter their function by substitution of modular subunits.

Studies of individual complexes^71^ as well as systematic investigations in yeast have suggested protein paralogs as mutually exclusive^98^. We therefore asked how prevalent paralogs, as defined by the EggNog database^99^, are in structurally consistent and mutually exclusive protein pairs. Figure 6J shows a substantial enrichment of paralogs in the mutually exclusive set where 201 out of 511 pairs (∼39%) are paralogs while only 9 out of 482 (∼2%) structurally consistent are paralogs. This suggests gene duplication is a common way for complexes to alter their functions through the subfunctionalization of paralogs.

To further investigate how cells have adapted to the scenario of multiple subunits competing for the same interface, we asked if subunits have different expression patterns. We therefore compared ProteomeHD.2 scores of mutually exclusive protein pairs to protein pairs that are structurally consistent. Figure 6K shows mutually exclusive pairs are more likely to have lower ProteomeHD.2 covariation values than structurally consistent proteins. We additionally observe lower RNA expression correlation (Fig 6L) and higher tissue specificity (Fig S6B) among mutually exclusive subunits relative to structurally consistent pairs. We also explored subcellular localization as an alternative regulator of mutual exclusivity. We compared subcellular localization annotations from HPA^84^ of mutually exclusive pairs as well as annotations of structurally consistent pairs (Fig S6C). We observe no statistical difference between the two sets. Interestingly, we do see examples of mutually exclusive pairs in separate localizations (see ANGEL1 and EIF4EBP1 above), however this appears rare. Taken together, this suggests that mutually exclusive subunits are regulated primarily at the level of expression rather than subcellular localization.

### Lineage - specific expression of protein complexes and mutually exclusive subunits

Given the critical role of expression regulation in shaping complexes with specialized functions through mutually exclusive subunits, we next explored the potential implications of context-specific expression of these subunits in cancer lineages. To investigate this, we turned to a recent large-scale study that reported proteomes for 949 human cancer cell lines, derived from 28 tissues and over 40 different cancer types^100^. We first examined whether hu.MAP3.0 complexes exhibited differential expression between cancer lineages and identified complexes whose expression most varies across the different cancer lineages (Fig S7). Among them, the MCM complex (huMAP3_00154.1), which is upregulated in blood cancer lineages, including leukemias and lymphomas (Fig 7A, Fig S8). Although from this dataset alone we cannot determine whether MCM upregulation reflects disease progression, upregulation of the MCM complex has previously been associated with poor overall survival in lymphoma^101^.

**Figure 7.**
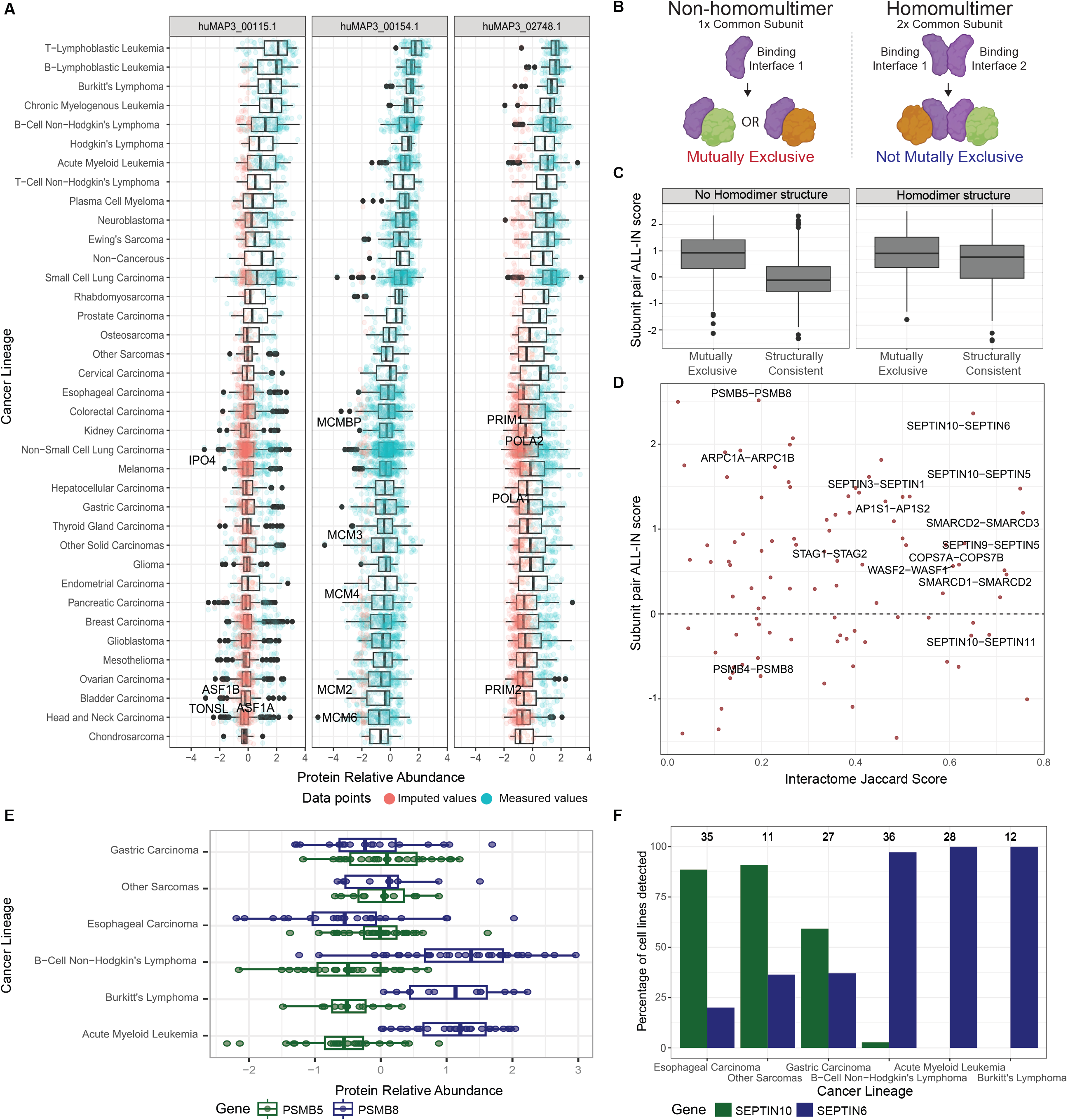
**Lineage - specific expression of mutually exclusive subunits.** (**A**) Relative protein expression across cancer lineages for three hu.MAP3.0 complexes that exhibit a high expression in leukemia lineages. Each point represents one subunit in one cell line. Where a subunit was not detected in a particular cell line, a low abundance value has been imputed. (**B**) Schematic of how a common subunit multimerization state affects mutually exclusive subunits (created with Biorender). Left side shows a common subunit (purple) that does not homomultimerize. This common subunit has only one binding site for two mutually exclusive subunits (green and orange). Right side shows a common subunit that homomultimerizes to allow for two binding sites, which accommodate two binding subunits (green and orange). (**C**) Boxplot of the expression-based mutual exclusivity score (ALL-IN) for structurally consistent and mutually exclusive interactions, respectively. Mutually exclusive subunits have a significantly higher ALL-IN score, and this effect is more pronounced for interactions that do not involve a homomultimerizing common subunit (Welch’s *T*-test for independent samples, \*\*\*\**P* <0.0001; box corresponds to quartiles of the distribution, line in box corresponds to median, whiskers extend to the first data point inside the first/third quartile plus 1.5 × interquartile range (IQR), respectively). (**D**) Subunit pairs that were predicted to be mutually exclusive by structural models are separated by their hu.MAP3.0 interactome overlap along the x-axis, and by their expression-based mutual exclusivity score (ALL-IN) along the y-axis. For a pair on the top right, the ALL-IN score can be seen as providing firm support for structure-predicted mutual exclusivity, and the pair shares many interaction partners in common, e.g. core subunits. Subunit pairs involving homomultimerizing common subunits were excluded from this plot. (**E**) Boxplots of relative protein abundance levels across the top 3 lineages that exhibit low and high expression of PSMB5 and PSMB8, respectively. (**F**) Percentage of cell lines per cancer lineage that express the SEPTIN10 and SEPTIN6 proteins at detectable levels. Number of cell lines per lineage are shown above the bars.

While our analysis revealed that cancer lineages vary significantly in terms of how many complexes are differentially expressed (Fig S8), most hu.MAP3.0 complexes do not show lineage-specific expression variation as a unit (Fig S7). We therefore assessed whether individual subunits of these complexes may be expressed in a lineage-specific manner.

Specifically, we focussed on potential mutually exclusive subunits. For all subunits of hu.MAP3.0 complexes we calculated a pairwise mutual exclusivity score, which we named ALL-IN (<u>A</u>verage cel<u>L L</u>ineage co-expressIo<u>N</u>) score, as it integrates the following four expression-based features (see methods): (1) correlation of protein abundances across the 949 cancer cell lines; (2) correlations aggregated by lineage; (3) a metric capturing very large expression differences (expression vs non-expression); (4) correlation with other subunits of the complex. To evaluate, we divide the mutually exclusive pairs into two groups based on whether the common subunit is known to form homomultimers (see methods). We anticipate common subunits that homomultimerize provide additional binding sites that can accommodate additional subunits alleviating the conflict of mutually exclusive subunits (Fig 7B). Indeed, we see pairs with common subunits that are non-homomultimers are much more likely to have a high mutually exclusivity score as opposed to subunits associated with common subunits that homomultimerize. This suggests pairs associated with common subunits that are non-homomultimers are more likely mutually exclusive (Fig 7C). For these protein pairs, we thus provide multiple lines of evidence to support their function as mutually exclusive complex subunits: shared hu.MAP3.0 complex membership, overlapping structural models, and divergent expression in cancer lineages (Fig 7D). Furthermore, for pairs that are covered by the cancer cell lineage dataset, it may be possible to link context-specific versions of protein complexes to different cancer lineages. For example, the PMSB8 subunit of the immunoproteasome is more abundant in blood lineages than PSMB5, the canonical proteasome subunit it replaces in these cell types (Fig 7E). Similarly, we identified SEPTIN6 and SEPTIN10 as mutually exclusive interactors of SEPTIN5 (Fig 6I, 7D). It has been previously noted that Septins are distributed differently across cell types, which could suggest expression as a potential regulator of these mutually exclusive interactions^95^. Indeed, we observe that SEPTIN6 is expressed in blood lineages, but is not detectable in lineages where SEPTIN10 is detected, such as gastric and esophageal lineages (Fig 7F). Importantly we demonstrate how the ALL-IN score can capture both protein pairs that anti-correlate in their expression patterns (Fig 7E), but also those that are not expressed in specific lineages highlighting their mutual exclusivity (Fig 7F).

## Discussion

Here we describe our integration of >25,000 proteomic experiments to build a more complete and accurate set of protein complexes. We advance on our previous work in several aspects. First, we place 13,769 human proteins into at least one multi-subunit protein complex. This is an increase of ∼40% over hu.MAP2.0. Second, we evaluate our protein complexes with respect to the protein co-expression network, ProteomeHD.2. Protein co-expression networks demonstrate considerable power in the identification of protein functional modules. Since protein co-expression networks are compiled from separate datasets than datasets used in our construction of a human protein complex map, interactions that agree between the two networks are considered very confident as there are multiple independent lines of evidence for their existence. Third, we developed a workflow to identify mutually exclusive pairs of proteins that likely do not co-exist in a complex based on structural modeling. This set of protein pairs point to potential complexes with multiple functions. As structural models of protein interactions become more widespread, we anticipate to find many more mutually exclusive pairs in protein complexes. Finally, we develop the ALL-IN score which is a prediction score of mutual exclusivity for protein pairs based on protein interactions and expression data. This is a general approach that can be applied to new interaction networks and expression datasets as they become available.

Mutually exclusive interactions introduce a conflict for organisms to resolve as two proteins are in competition for a single interface. We observed the dominant approach for organisms to relieve this conflict is to alter the expression of mutually exclusive pairs so they are not present concurrently in the same cell (Fig 6K,L). Other possible mechanisms for organisms to relieve this conflict include altering the subcellular localizations of mutually exclusive pairs and regulating protein subunits at the level of post translational modifications (PTMs). We see limited evidence for mutually exclusive pairs having altered their subcellular localization compared to structurally consistent pairs (Fig S6C). This is in line with other groups who have explored whether paralogs in yeast have differing subcellular localization and found duplicated genes are not more likely to relocalize compared to singleton genes^102^. Post translational modification of individual mutually exclusive subunits is another potential mechanism to activate or deactivate a protein subunit’s membership in a complex. Many interactions are known to be modulated by PTMs^103^. Although we don’t expect PTMs to play as large of a role as expression, we do anticipate exploring this mechanism further in the future.

Another possibility to relieve the conflict of mutually exclusive pairs is protein multimerization. In this work we make the assumption that there is one to one stoichiometry between the common protein’s binding site and a mutually exclusive subunit. However, the cell may relieve the conflict by having multiple binding sites (e.g. multimerize the common protein) to accommodate the two mutually exclusive proteins. For example, proteins that form fibers such as actin can support many proteins binding to the same interface yet at different protein instances along the fiber. In Figure 7B,C, we explore the possibility of multimerization alleviating mutual exclusive conflicts. We evaluate common proteins that are found to homo-multimerize from those that do not and see those that do not homo-multimerize are more likely to have mutually exclusive binding sites. This suggests multimerization is another avenue for organisms to address the conflict of mutual exclusive interactions.

Also implicit in our definition of mutual exclusive pairs is that the pairs are required to sterically hinder each other. An alternative class of mutually exclusive pairs may include those that allosterically alter the common protein surface to restrict the binding of the other protein.

Additional modeling of alternative conformations is likely required to identify this alternative class. We also make sensible cut-offs such as quality of models and number of overlapping interface residues to identify mutually exclusive pairs. We therefore anticipate as structural modeling of protein interactions becomes more accurate, we will identify additional mutually exclusive interactions.

Finally, we imported the 15,326 hu.MAP3.0 clusters into the Complex Portal and matched at the canonical level to existing manually curated entries. No thresholds were imposed in the minimum number of participants required to make the match. Where the protein components are an exact match to an existing manually curated complex, the entries have been merged and the hu.MAP3.0 accession number retained as a cross-reference in the entry, with the qualifier ‘identity’. Where the hu.MAP complex is a subset of an existing manually curated entry, or entries, again these have been merged and the hu.MAP3.0 cross-reference added, with the qualifier ‘subset’. The remaining complexes were created as novel assemblies into the Complex Portal^21^ and have been publicly released for search and download. The complexes will be priority targets for manual curation based on literature curation and comparison to small-scale interaction datasets in the IMEx Consortium dataset^104^, and to other curated datasets in resources such as PDB^105^, EMDB^106^ and Reactome^107^. The use of LLMs to enable rapid curation of these assemblies based on the presence of sets of gene names in a publication is also being evaluated. Overall, we anticipate this to be an extremely valuable tool for the broad research community.

## Methods

### Reagent and Tools table

| Reagent/Resource | Reference or Source |
| --- | --- |
| MCL Version: 14-137 | https://micans.org/mcl/106 <sup>26</sup> |
| ClusterOne Version: 1.0 | https://paccanarolab.org/clusterone/ <sup>25</sup> |
| ProteinComplexMaps Version: bdea06b | https://github.com/KDrewLab/protein_complex_maps <sup>16</sup> |
| AutoGluon Version 0.4.0 | https://auto.gluon.ai/stable/install.html <sup>24</sup> |
| Training of hu.MAP 3.0 with AutoGluon | https://github.com/KDrewLab/huMAP3.0_analysis |
| EvalMutualExclusivity | https://github.com/KDrewLab/huMAP3.0_analysis |
| ~10,000 AlphaFold2 structures for mutually exclusive dimer modeling | https://archive.bioinfo.se/huintaf2/humap.zip <sup>108</sup> |

### Curation of mass spectrometry datasets

To build the hu.MAP 3.0 network, we integrated data from more than 25,000 previously published mass spectrometry experiments. Leveraging our specialized machine learning framework, we enhanced our previously published hu.MAP 2.0 network by incorporating 10,000 new affinity purification data from Bioplex 3.0^1^. Bioplex 3.0 expands ORFeome coverage with over 10,000 baits in HEK293T cells and biological diversity by including HCT116 cells, significantly enriching the interactome within our network.

To construct a machine learning classifier, we collected mass spectrometry data from multiple publications and derived features predictive of protein interactions. Protein interaction features for datasets used in hu.MAP 2.0^16^ were downloaded from http://humap2.proteincomplexes.org/static/downloads/humap2/humap2_feature_matrix_20200820.featmat.gz. This matrix, containing 292 features, has been previously described^16^. Briefly, the features were obtained from original sources or calculated directly from raw mass spectrometry data. Of these, 220 features are derived from 55 CF-MS datasets^8^ and include metrics such as Poisson noise Pearson correlation coefficient, weighted cross-correlation, co-apex score, and MS1 ion intensity distance. Twenty-nine features are sourced from AP-MS data, including HGScore^109^, MEMOs (core modules) certainty assignments^7^, and metrics from Bioplex 1.0^3^ and 2.0^2^, such as NWD Score, *Z* Score, Plate *Z* Score, Entropy, Unique Peptide Bins, Ratio, Total PSMs, Ratio Total PSMs, and Unique:Total Peptide Ratio. Additional AP-MS features include prey.bait.correlation, log10.prey.bait.ratio, og10.prey.bait.expression.ratio, mean psm^4^, and socioaffinity index terms^5^. Fifteen features come from proximity labeling datasets^14,15^, including metrics Average Spectra, Average Saint probability, Max Saint probability, Fold Change, and Bayesian FDR estimate. Data sources for datasets incorporated into hu.MAP3.0 are highlighted in Table 1.

**Table 1:** MS Datasets incorporated into hu.MAP 3.0

| Dataset | Number of Experiments | Link |
| --- | --- | --- |
| Wan et al (2015) | 5,344 fractions | http://hu1.proteincomplexes.org/static/downloads/feature_matrix.txt.gz |
| Hein et al (2015) | 1,125 bait pull-downs | http://hu1.proteincomplexes.org/static/downloads/feature_matrix.txt.gz |
| Gupta et al (2015) | 2 conditions x 58 proximity labeled baits = 116 | http://prohits-web.lunenfeld.ca/ |
| Youn et al (2018) | 119 proximity labeled baits | http://www.cell.com/cms/attachment/2118963855/2087347233/mmc2.xlsx |
| Boldt et al (2016) | 217 bait pull-downs | https://static-content.springer.com/esm/art%3A10.1038%2Fncomms11491/MediaObjects/41467_2016_BFncomms11491_MOESM835_ESM.xlsx |
| Treiber et al (2017)<br>(reprocessed by (Mallam et al, 2019)) | 3,004 RNA hairpin pull-down MS runs | https://www.cell.com/cms/10.1016/j.celrep.2019.09.060/attachment/45abb95b-ef3f-4752-8906-dc5eed118480/mmc4.csv |
| Bioplex 3.0 (Huttlin et al 2021 )<br>(includes Bioplex 2.0 and 1.0) | 10,128 bait pull-down HEK 293T, 5,522 bait pull-downs HCT 116 | https://bioplex.hms.harvard.edu/data/BioPlex_BaitPreyPairs_noFilters_293T_10K_Dec_2019.tsv,<br>https://bioplex.hms.harvard.edu/data/BioPlex_BaitPreyPairs_noFilters_HCT116_5.5K_Dec_2019.tsv |

Newly integrated Bioplex 3.0 data sets comprised the same measures that were used for Bioplex 2.0 and Bioplex 1.0 features obtained (see Table 1), including specifically NWD Score, *Z* Score, Plate *Z* Score, Entropy, Unique Peptide Bins, Ratio, Total PSMs, Ratio Total PSMs, Unique:Total Peptide Ratio and Average Assembled Peptide Spectral Matches. The features are obtained from both the HEK293T and HCT-116 datasets.

Additional features were generated using a weighted matrix model (WMM), which is based on the hypergeometric distribution as described in^23^ and ^81^, and previously used in ^16^. WMM features are derived from the presence or absence of proteins across individual experiments. To mitigate the effects of noise common in high-throughput mass spectrometry data — such as the spurious identification of proteins with low counts in single experiments — we applied thresholds to ensure that only high-quality identifications were considered. The datasets utilized were filtered according to the protocols established in^16^. We calculated WMM features for each BioPlex 3.0 cell type as well as a combined set (i.e. HEK293T, HCT-116, HEK293T/HCT-116).

Additionally, we calculated WMM features for two abundance thresholds, where a given protein was required to have > 2.0 Bioplex3.0 *Z* score and > 4.0 Bioplex3.0 *Z* score. For each calculation, we generate a feature in the form of the negative natural log P-value of Equation (1) and the total number of experiments the pair of proteins is observed together (i.e., pair count).

The final feature matrix contains 324 features for 25,992,008 protein pairs.

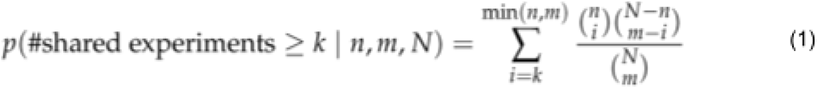

### Gold standard protein complex test and train sets

To generate a test and training set of protein complexes, we downloaded the complete set of human protein complexes from the Complex Portal^110^, version 2022_07_11 (https://ftp.ebi.ac.uk/pub/databases/intact/complex/2022-07-11/). The Complex Portal is a manually curated database of stable protein complexes. Isoforms were removed. Complexes were randomly assigned to the test or train set. To prevent overlap between these sets, any overlapping complexes (i.e., those sharing protein pairs) were identified and the largest complex was removed, ensuring that no PPI was present in both sets. Additionally, complexes greater than 30 subunits were removed to prevent large protein complexes from dominating the test and training sets. Within the test and train sets, pairs of proteins were generated and labeled as positive and negative. Positive pairs are proteins within the same complex; negative pairs are proteins in separate complexes. Any intersecting PPIs between the test and training sets were identified and removed to prevent overlap. The split_complexes.py script in the python package protein_complex_maps (Reagents and Tools Table) was used for test and training set generation. Below are annotated command-line examples used to reduce redundancy in the gold standard complex lists and split complexes into test and training sets:

### Reduce redundancy of benchmark complexes

To remove redundancy within benchmark complexes, the benchmark complex file was formatted as tab separated lists of UniProt identifiers, with one complex per line. We then ran the python script `complex_merge.py` using the –remove_largest flag which removes the largest of redundant pairs of complexes. The input file formatted from Complex Portal can be found in Table S5.

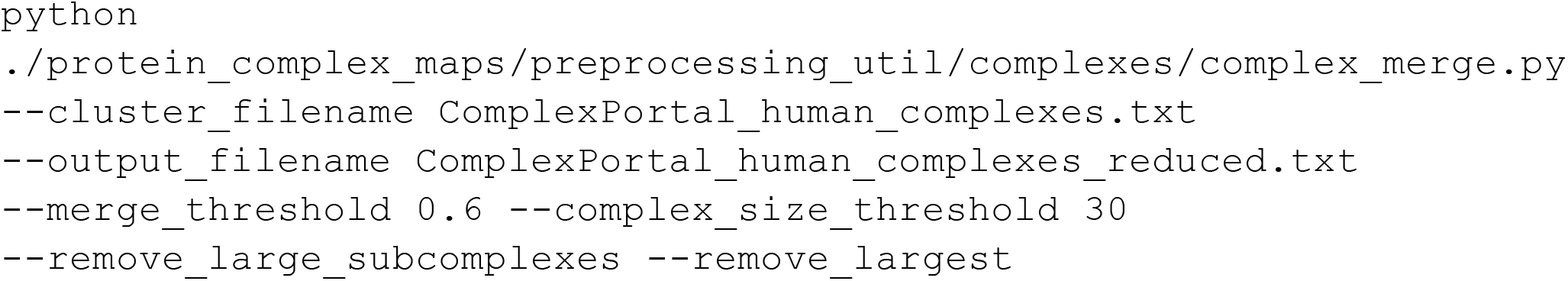

### Generating test and train sets

The resultant redundancy reduced complex file can then be used as an input for `split_complexes.py`, which generates six output files: negative training PPIs (input_filename.neg_train_ppis.txt), negative test PPIs (input_filename.neg_test_ppis.txt), positive training PPIs (input_filename.train_ppis.txt), positive test PPIs (input_filename.test_ppis.txt), the complexes used for training PPI generation (input_filename.train.txt), and the complexes used for test PPI generation (input_filename.test.txt). All six resultant files used in this analysis can be found in Table S5.

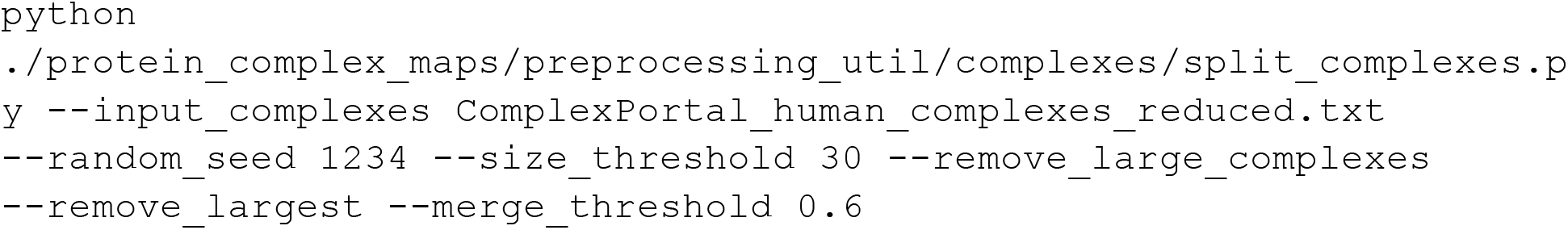

### Model Description

To generate a protein interaction network, we utilized AutoGluon^111^, an open-source automated machine learning framework, to classify pairs of proteins as co-complex interactions. AutoGluon efficiently explores a broad range of model architectures, selecting those that best optimize the predictive accuracy of the final model.

To train the model, we constructed a feature matrix with protein pairs as rows and features as columns. Positive and negative labels were assigned using the benchmark training set. We randomly subsampled 10,000 negative labeled pairs to balance positive and negative labels. We used the AutoGluon TabularPredictor module using the constructed feature matrix to fit a model that discriminates between positive and negative interactions. We evaluated three different evaluation metrics, including accuracy, F1, and precision. The final model was selected based on its AUPRC (area under the precision-recall curve) on the benchmark leave-out test set, with accuracy optimization yielding the top-performing model (Fig S1A).

During training, AutoGluon evaluated 13 different base models, and a final ensemble model. The final ensemble model consisting solely of XGBoost with a weight of 1, was the best performer (Table S6). We used this final model on all protein pairs within the feature matrix to predict whether they are interactions or not, including a confidence score. Links to Jupyter notebooks containing modeling training, testing, and prediction generation are provided in the Reagents and Tools Table.

We evaluated the final model using precision-recall analysis on the benchmark leave-out test set and compared its performance to other datasets, including our previous hu.MAP 2.0 model^16^ and BioPlex 3.0^1^ (Fig 2A). All pairwise predictions for protein pairs in hu.MAP 3.0 are available in Table S7.

### Clustering and parameter set selection

To identify clusters within the protein interaction network we used a two-staged clustering approach outlined above (Fig 2B). We first threshold the network based on the confidence score output from the model. To the thresholded network we applied the ClusterOne^25^ algorithm to identify dense regions within the network. For each identified dense region, we applied the MCL^26^ algorithm to further identify clusters. To determine optimal parameters for cluster identification, we ran parameter sweeps generating clusters with various parameter combinations. Specifically, we used a range of parameters for model confidence score (0.95, 0.9, 0.85, 0.8, 0.7, 0.6, 0.5, 0.4, 0.3, 0.2, 0.1, 0.05, 0.04, 0.03), ClusterOne max overlap (0.6, 0.7), ClusterOne density (0.1, 0.2, 0.3, 0.4), and MCL inflation (1, 2, 3, 4, 5, 7, 9, 11, 15). In addition, after clustering we applied a filter to remove proteins from the resulting clusters which do not have edge weights greater than the model confidence score threshold, which infrequently arose as a result of the MCL algorithm. Finally, we only consider clusters where the number of subunits is <=100 proteins as larger clusters appeared, after manual inspection, to have disparate biological functions.

For clustering evaluation (Fig 2C,D), we used the *k*-cliques method, focusing specifically on weighted precision (P_weighted) and weighted recall (R_weighted) as detailed in our previous work^16,17^. In essence, the *k*-cliques method globally compares clusters to gold standard complexes by evaluating cliques — fully connected subgroups — extracted from the clusters against those from the gold standard complexes. This comparison spans all clique sizes from pairs (size 2) to the largest complex or cluster (size n). For each clique size, precision and recall are computed. These metrics are then averaged across all clique sizes, with weights assigned based on the number of clusters with size >= to the clique size, to reduce the influence of larger clusters on the overall precision and recall.

All clusterings were analyzed using the *k*-cliques method, with comparisons to the training set of gold standard protein complexes. From this, we manually selected six cluster sets that provided optimal balance between precision and recall, as illustrated in Figure 2C. These five clusterings were merged into a union set, and the parameters for these clusters are shown in Table 2.

**Table 2:** Clustering parameters

| Clustering | Confidence | Weighted clique precision | Weighted clique recall | F1 score weighted clique | Total clusters | Total proteins | Confidence score threshold | ClusterOne density | ClusterOne overlap | MCL inflation |
| --- | --- | --- | --- | --- | --- | --- | --- | --- | --- | --- |
| 1 | Extremely high | 0.72 | 0.31 | 0.43 | 5468 | 11221 | 0.95 | 0.2 | 0.7 | 15 |
| 2 | Very high | 0.66 | 0.34 | 0.45 | 5678 | 12049 | 0.95 | 0.3 | 0.7 | 5 |
| 3 | High | 0.63 | 0.39 | 0.48 | 5302 | 11970 | 0.95 | 0.1 | 0.7 | 4 |
| 4 | Moderately high | 0.58 | 0.44 | 0.5 | 5237 | 12346 | 0.95 | 0.1 | 0.7 | 3 |
| 5 | Medium high | 0.53 | 0.48 | 0.5 | 4827 | 12449 | 0.95 | 0.2 | 0.7 | 2 |
| 6 | Medium | 0.47 | 0.53 | 0.5 | 5503 | 13532 | 0.8 | 0.1 | 0.7 | 2 |

Final evaluation of the selected clusters, we used the *k*-cliques method^16^ against the gold standard leave-out test set complexes for all individual cluster sets and the union set (Fig 2D). We additionally compare against previously published Bioplex 3.0 protein complexes. The complete list of hu.MAP 3.0 complexes with UniProt identifiers and confidence scores are available in Table S1.

### Annotation enrichment analysis for hu.MAP 3.0 complexes

For all complexes identified in hu.MAP 3.0, we calculated annotation enrichment for GO, Reactome, CORUM, KEGG, and Human Phenotype Ontology (HP) using gProfiler^112^ (Table S2). We evaluate complex enrichment by comparing the results against a set of randomly shuffled complexes. In shuffled complexes, protein IDs were randomly reassigned to cluster IDs, maintaining the same size and distribution of clusters while preserving the original protein-per-complex distribution. We applied a p-value threshold of 0.01 for annotations in all categories. Additionally, we excluded annotations derived via electronic transfer and used the entire set of proteins from the ∼25,000 mass spectrometry experiments as the background.

Within this analysis, we also sought to identify uncharacterized proteins within the predicted hu.MAP 3.0 complexes to facilitate annotation transfer. We used UniProt annotation scores (www.uniprot.org/help/annotation_score), which range from 1 (least characterized) to 5 (most characterized), to classify proteins within predicted complexes and to identify uncharacterized proteins. To transfer annotations to previously uncharacterized proteins, complexes were required to meet the following criteria: (1) 50% of complex members must share the same annotation, (2) the annotation must have a corrected p-value <= 0.01, (3) the uncharacterized protein must have a UniProt annotation score <=3. Annotations were considered from KEGG, GO, CORUM, and Reactome. “CORUM root”, “REACTOME root term”, and “KEGG root term” were filtered out prior to analysis.

### Covariation analysis with ProteomeHD.2

ProteomeHD.2 is the second version of ProteomeHD, our previously reported protein covariation map^19^. ProteomeHD.2 will be described in detail in a separate manuscript (Kourtis et al, in preparation). In brief, the workflow for producing ProteomeHD.2 was similar to the one for ProteomeHD, involving the re-processing of 23,000 previously published mass spectrometry raw files deposited in the PRIDE repository^113^. Together, these data covered protein abundance changes in response to 2,498 biological perturbations (e.g. drug vs control treatments), all quantified using stable-isotope labeling by amino acids in cell culture (SILAC). To identify proteins with correlated abundance changes across these 2,498 conditions, an unsupervised covariation network was constructed using the treeClust algorithm^19^. In contrast to ProteomeHD, for ProteomeHD.2 an additional supervised machine-learning step was included: the covariation scores, together with a range of edge quality features (e.g. the number of peptides detected per protein or the number of shared samples in which both proteins were detected) were used to train a Random Forest model to distinguish between genuine protein-protein covariation and likely false positive interactions. A combination of protein pairs, known to be functionally related according to STRING, REACTOME or BIOGRID, were used as training data. The resulting Random Forest probabilities were used as final protein covariation score, and were integrated with the hu.MAP3.0 protein-protein interaction scores at the protein-pair level for downstream analysis.

### Structural Modelling of Uncharacterized Proteins

AlphaFold3 web server^56^ was used to model the three dimensional structure of protein interactions in Figure 4 & 5. Canonical sequences from UniProt were used for each protein. All defaults were used. TMEM167A and IER3IP1 are both identified as transmembrane proteins, therefore 12 copies of palmitic acid were included in the modeling.

### Identification and characterization of mutually exclusive protein subunits

To identify mutually exclusive interactions within hu.MAP3.0 complexes, we examined more than 10,000 AlphaFold2 structures of pairs of co-complex proteins^79^ (https://archive.bioinfo.se/huintaf2/humap.zip). We filtered AlphaFold2 structures using pDockQ score (> 0.23) as well as IF_pLDDT value (>70). A pDockQ score greater than 0.23 has been previously shown to correspond to 70% of structures being well-modeled for a single conformation^79^. Additionally, we ensured that AlphaFold2 models had <5 overlapping residues between protein chains.

To classify two proteins as 1) mutually exclusive or 2) structurally consistent, we identified AlphaFold2 models that contained a third shared protein subunit. We aligned these structures using the PyMol^114^ align function on the third shared subunit. To determine if the two proteins are mutually exclusive or structurally constituent, we compare their interfaces with the third protein for overlap. We define the interface of each protein with the third shared protein as atoms within 4.0 Å of the third shared protein. Interface residues between each protein were tested for overlap (residue atoms < 4.0 Å). Pairs with >10 overlapping interface residues were considered mutually exclusive. Pairs with 0 overlapping interface residues were considered structurally consistent. Due to expected error in AlphaFold2 models, we considered pairs with < 10 and > 0 overlapping interface residues inconclusive. We tracked residue overlap between unique protein chains, regardless of their presence at defined interfaces, to further filter the structurally consistent pairs. Overlapping could result from clashes in poorly modeled regions and does not necessarily indicate structural inconsistency. Therefore, we excluded structurally consistent pairs which had > 100 overlapping residues between their unique protein chains.

Link for code repository, used to determine mutually exclusive pairs, is available in the Reagents and Tools Table.

We calculated annotated enrichment for GO terms (e.g., GO:MF, GO:BP, and GO:CC) of mutually exclusive protein pairs using g:Profiler^112^. We took all proteins (i.e., the common and unique subunits) of identified mutually exclusive interactions (539 proteins) and compared them against all ∼1100 proteins identified between the mutually exclusive and structurally consistent groups as the background.

We analyzed potential causes of mutual exclusivity, focusing on: 1) Paralogs between mutually exclusive and structurally consistent protein pairs, 2) Covariation of these pairs, 3) Differential tissue expression, and 4) Subcellular compartmentalization. For paralog identification, we used eggNOG^99^ ortholog groups at the Eukaryota level and determined if both unique proteins in each pair belonged to the same ortholog group. Covariation was assessed using ProteomeHD.2 scores^38^, which reflect the probability that two proteins covary across cell types and conditions. To evaluate the expression similarity across tissues, we used RNA-seq data from Human Protein Atlas^84^ (https://www.proteinatlas.org/download/rna_tissue_consensus.tsv.zip) and calculated Pearson’s correlation coefficient of normalized RNA expression (nTPM) across tissues. We also analyzed tissue specificity scores (https://www.proteinatlas.org/about/assays+annotation#classification_rna) from the Human Protein Atlas^84^ and compared the number of pairs across categories (Fig S6B). Lastly, we compared subcellular localization using Jaccard similarity coefficients (0 = no similarity, 1 = identical) between the unique proteins, based on data from the Human Protein Atlas (https://www.proteinatlas.org/download/proteinatlas.tsv.zip) (Fig S6C). All of these metrics are annotated within the table of identified mutually exclusive pairs (Table S4).

### Cancer lineage-related protein expression analysis

Cancer lineage-specific protein expression data were extracted from supplementary table 2 (Protein matrix) of Gonçalves et al^100^. Log-scale protein intensities were z-scored for downstream analysis, thereby reflecting protein expression changes relative to their mean abundance across all cell lines. We refer to these z-scores as relative protein abundances. These data were used to create the ALL-IN (<u>A</u>verage cel<u>L L</u>ineage co-expressIo<u>N</u>) score, designed to reflect the likelihood of a protein pair to be mutually exclusive subunits of a protein complex. To create the ALL-IN score we first calculated four separate sub-scores for all protein pairs annotated as part of the same hu.MAP3.0 complex:

1. Correlation of protein abundance changes across the 949 cell lines, determined using robust correlation (bicor)^115^. Unlike the machine-learning based similarity metrics used for ProteomeHD, correlation metrics can detect anti-correlation (negative correlation), which is expected for mutually exclusive subunits.
2. Robust correlation of protein abundance changes after averaging protein intensities by cell lineage. By averaging values across lineages, a potential bias in subscore 1 stemming from different lineages having different numbers of cell lines, is removed.
3. A sub-score reflecting very large expression differences. Sub-scores 1 and 2 fail to detect very large expression differences, because it is not possible to obtain correlations for samples where a protein was below the detection threshold. To overcome this, we first subset the dataset to include proteins that were detected in at least 50 cell lines. The vast majority of proteins in the resulting dataset had z-scores between -2 and 2. We then impute missing values by random sampling from a normal distribution centered around -4 with a standard deviation of 0.3. Next, for each protein, we consider the 50 cell lines with the highest and lowest expression levels (including imputed values), respectively. For each protein pair, we then calculate the difference between the average expression level (z-score) of protein A in the top 50 and the bottom 50 cell lines of protein B, and vice versa. We combine the values for protein A and B by averaging. The resulting sub-score reflects the degree of differential protein expression between A and B. Unlike sub-scores 1 and 2 this takes into account missing (imputed) values, i.e. cell lines where one of the two proteins was not detectable.
4. For each subunit, we calculate the average robust correlation to all other subunits of a complex, separately for each lineage. This results in a correlation coefficient indicating how well each protein correlates with the rest of the complex in each lineage. For each protein pair we then correlate these lineage-specific correlations. The rationale of this sub-score is that, for mutually exclusive subunits expressed in a lineage-specific manner, the dominant subunit in a lineage should correlate well with the rest of the complex in that lineage. By identifying pairs which differ in their lineage-specific correlation with the main complex we can pinpoint possible mutually exclusive pairs.

To combine these four sub-scores into the ALL-IN score for mutually exclusivity, each sub-score was z-scored and the sub-scores averaged. Only protein pairs with at least two out of the four possible sub-scores were included. To address how mutual exclusivity might be influenced by the presence of target subunits existing in multimeric states—such as homodimerization of common protein subunits providing multiple equivalent binding sites (see Fig 7A,B)—we annotated whether the common protein is known to homomultimerize^116^ and compare the ALL-IN scores across these groups. Non-homodimeric mutually exclusive protein pairs and their subunit ALL-IN scores can be found in Table S8.

### Overexpression testing of hu.MAP.3 complexes in cancer lineages

Proteins belonging to confidence 1 hu.MAP.3 complexes with more than 3 subunits in the cancer lineage dataset, and with less than 80% missing values, were z-scored as indicated in the previous paragraph. The missing values were then imputed with a normal distribution of mean -2 and standard deviation of 0.3, followed by a second round of z-scoring. The subunit relative abundance scores were averaged to the complex-level per cell line and the enrich function from the EnrichIntersect R package^117^ was used to calculate the overrepresentation score and p-values, which were subsequently FDR-adjusted. Table S9 includes data of protein-level abundance, average complex abundance per cell line, complex expression by lineage, and enrichment scores by lineage.

## Data Availability

Both the manually curated set of reference protein complexes and hu.MAP3.0 predicted assemblies are available from the Complex Portal. Data can be accessed either via the website (www.ebi.ac.uk/complexportal), our ftp site (ftp.ebi.ac.uk/pub/databases/intact/complex/current/) or our REST API (https://www.ebi.ac.uk/intact/complex-ws/). The FTP site contains files in different formats, grouped by species, and for each species there are separate files containing manually curated complexes and ML-predicted complexes. The Complex Portal is an open source (Apache 2.0), open data (CC0) project, details on https://www.ebi.ac.uk/complexportal/about#license_privacy . Complexes can also be searched and downloaded using the hu.MAP3.0 website (https://humap3.proteincomplexes.org). The complete hu.MAP3.0 complex map, protein interactions with ML confidence scores, test and training data, and feature matrix can be found on https://humap3.proteincomplexes.org/download . Software used to generate and analyze the hu.MAP3.0 complex map can be found on GitHub (https://github.com/KDrewLab/protein_complex_maps). Software used to train, test, and make predictions with huMAP3.0 can be found on GitHub (https://github.com/KDrewLab/huMAP3.0_analysis). Software used to identify mutually exclusive protein pairs and analyze their features can be found on GitHub (https://github.com/KDrewLab/huMAP3.0_analysis). Software related to ProHD2 and ProCAN covariation, and their integration with hu.MAP3.0 can be found on GitHub (https://github.com/Skourtis/ProHD2_huMAP3). All pairwise ProteomeHD.2 scores are available upon request.

## Author Contributions

SNF, ERC, SK, SO, HH, GK, and KD conceived of the study. SNF, ERC, SK, GK, and KD designed experiments and analyzed data. SNF, ERC, SK, GK, and KD wrote software including software pipeline and web resources. SNF, ERC, SK, SS, SO, HH, GK, and KD discussed and interpreted the results, and wrote the manuscript.

## Acknowledgements

This work was supported by grants from the BBSRC - National Science Foundation/Directorate for Biological Sciences [BB/X002179/1][BB/X002683/1], National Human Genome Research Institute (NHGRI), Office of Director [OD/DPCPSI/ODSS]; National Institute of Allergy and Infectious Diseases (NIAID), National Institute on Aging (NIA), National Institute of General Medical Sciences (NIGMS), National Institute of Diabetes and Digestive and Kidney Diseases (NIDDK), National Eye Institute (NEI), National Cancer Institute (NCI), National Heart, Lung, and Blood Institute (NHLBI) of the National Institutes of Health [U24HG007822] and European Molecular Biology Laboratory (EMBL) core funds. GK is supported by an MRC Career Development Award (MR/T03050X/1) and a Royal Society Research Grant (RGS\R2\212303). KD and SNF are supported by National Institute of Child Health and Human Development (R00 HD092613). We thank Eliot Ragueneau for artwork design.

## Tables

Reagents and tools
Table 1: MS datasets incorporated into hu.MAP 3.0
Table 2: Summary of key clustering parameters
Table S1: Complete list of hu.MAP3.0 complexes and confidence scores
Table S2: GO enrichments of hu.MAP3.0 complexes
Table S3: Annotations for uncharacterized proteins in hu.MAP3.0 complexes
Table S4: Mutually exclusive protein pairs and metrics
Table S5: Training and test sets
Table S6: Performance metrics for AutoGluon models
Table S7: Predicted pairwise probability with associated hu.MAP3.0 complexes
Table S8: ALL-IN scores for nonhomodimeric mutually exclusive protein pairs
Table S9: ProCAN expression data and scores

---

**Figure S1.**
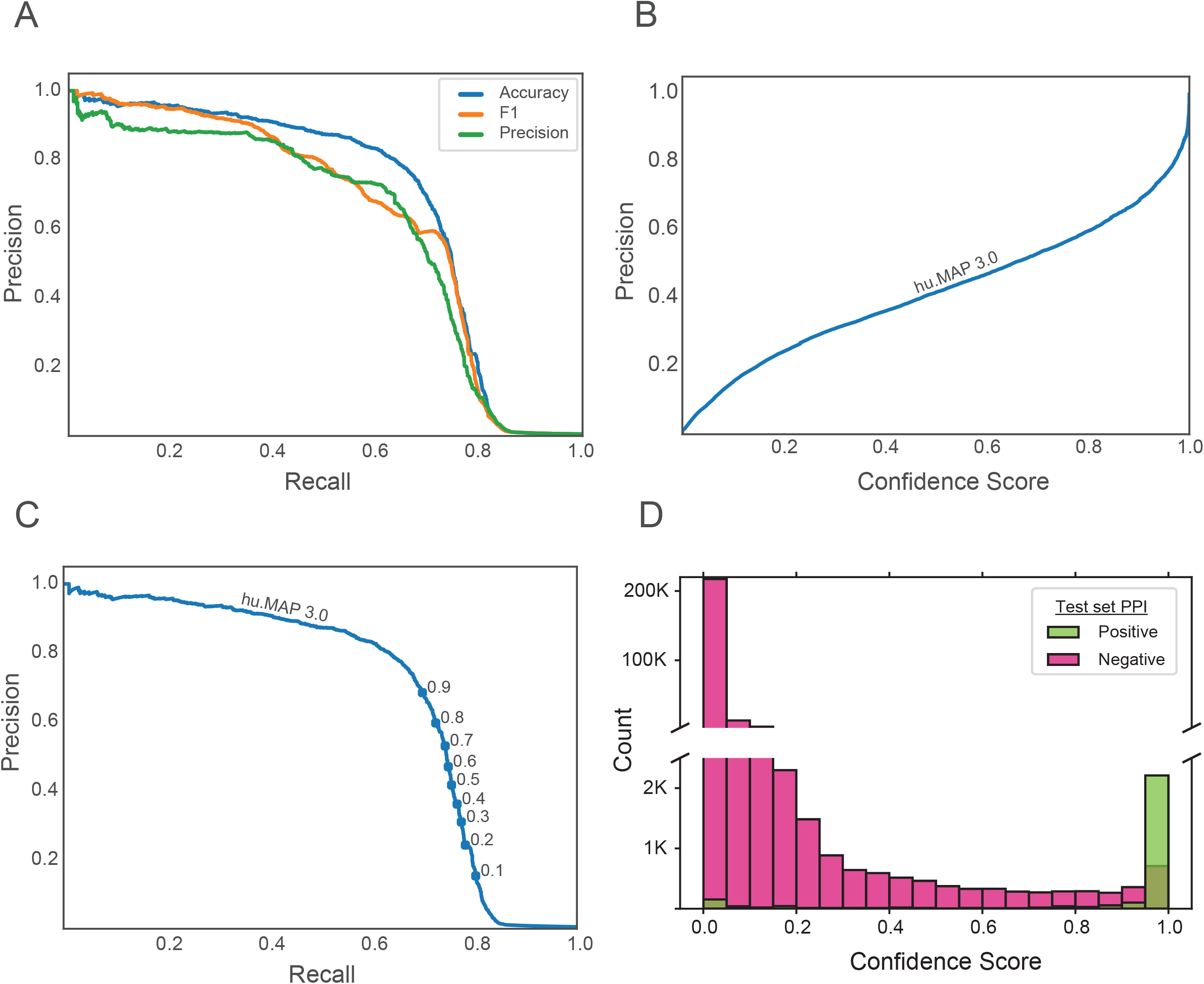
Characterization of hu.MAP 3.0 model performance. (**A**) Precision-Recall plot evaluated on the leave-out test set of gold standard interactions for different versions of hu.MAP 3.0. The different models were trained with the same literature-curated training set but optimized for different evaluation metrics. The accuracy optimized model shows the greatest performance and is designated as the hu.MAP 3.0 model. (**B**) Line plot depicting the relationship between precision and the hu.MAP 3.0 model confidence score on the test set interactions, illustrating that increases in confidence score correspond with increased precision. (**C**) Precision-Recall plot for the hu.MAP 3.0 model on test set interactions, with confidence score thresholds marked as circles, demonstrating clear separation along the curve. (**D**) Distribution of positive and negative test set interactions across model confidence scores. The histogram highlights the separation of these interactions.

**Figure S2.**
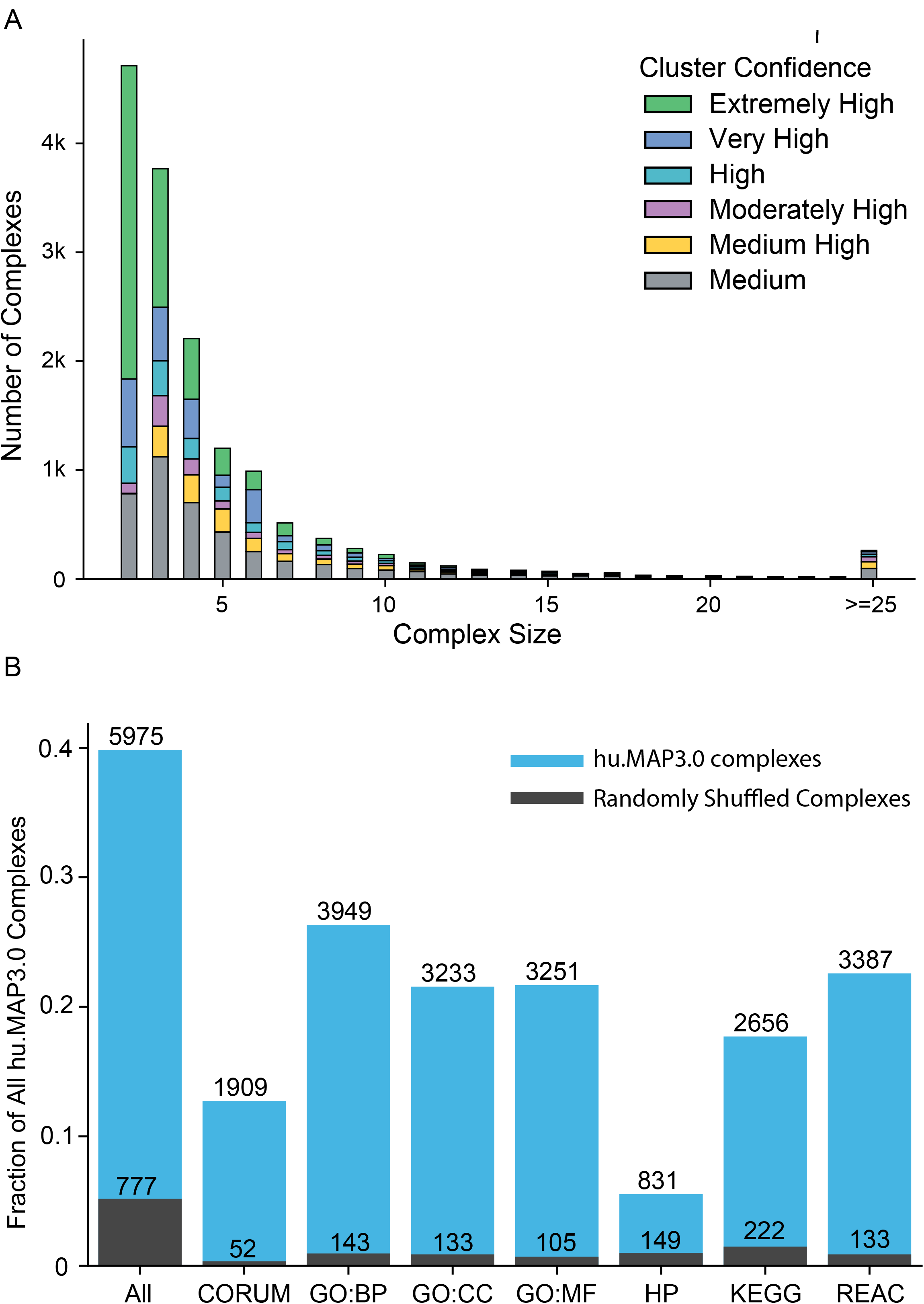
Clustering size distribution and annotation enrichment. (**A**) Distribution of hu.MAP3.0 complex size. Colors in stacked bars represent cluster confidence. (**B**) Annotation enrichment of hu.MAP3.0 complexes using g:Profiler. Bars represent number of complexes enriched for GO, KEGG, CORUM, Reactome, or Human Phenotype Ontology (HP) annotations that pass a corrected p-value of 0.01. Random represents enriched protein sets when complex membership is shuffled for all complexes.

**Figure S3.**
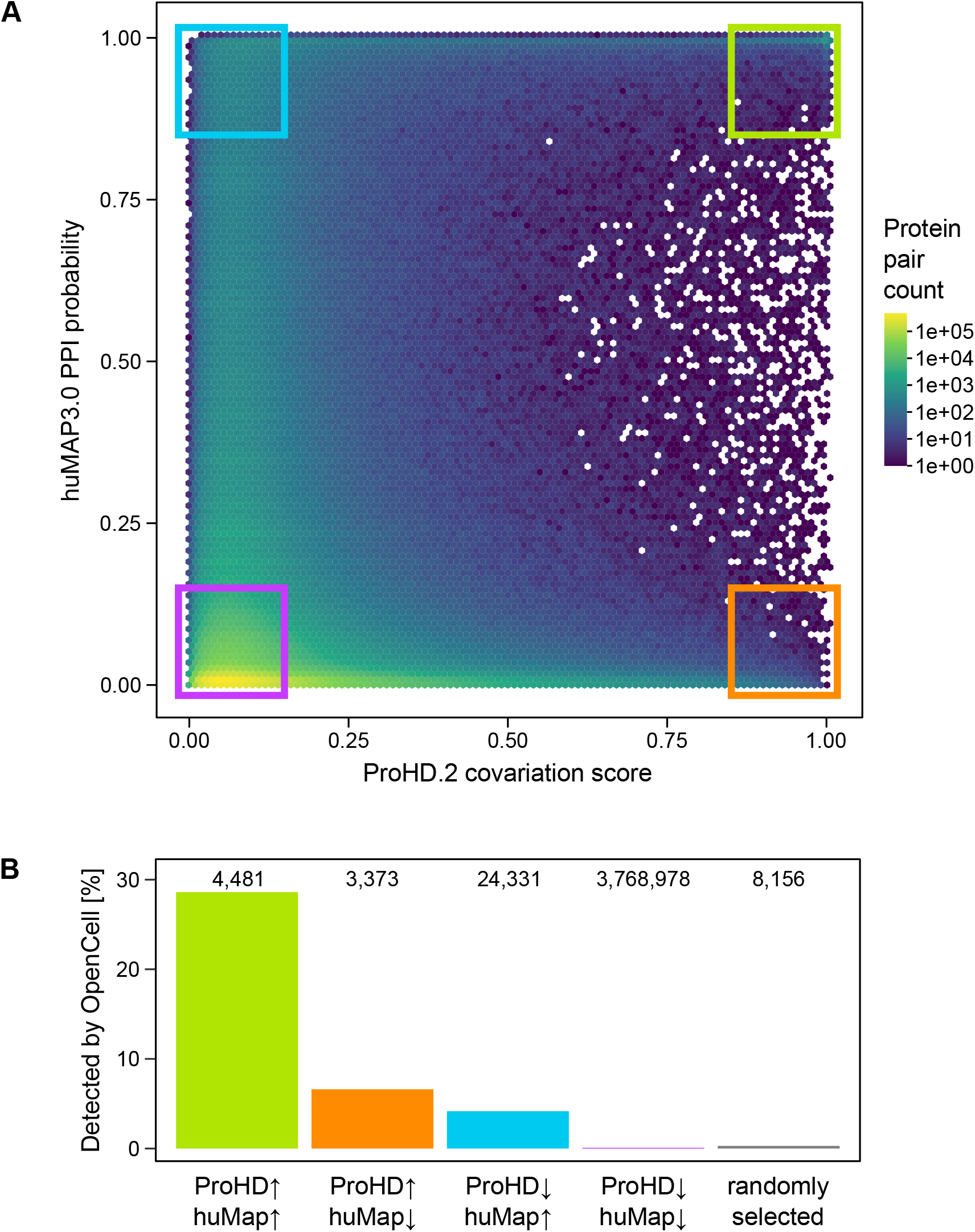
Comparison of ProteomeHD.2, hu.MAP3.0, and OpenCell. (**A**) Scatterplot of protein pairs’ ProteomeHD.2 covariation score (x-axis) and hu.MAP3.0 confidence score (y-axis). Four quadrants are highlighted where yellow = high in hu.MAP3.0, high in ProteomeHD.2, blue = high in hu.MAP3.0, low in ProteomeHD.2, orange = low in hu.MAP3.0, high in ProteomeHD.2, and purple = low in both. (**B**) Comparison of four quadrants in (**A**) to an orthogonal protein interaction dataset, OpenCell, not included in either hu.MAP3.0 nor ProteomeHD.2. Bars represent the percent detected in OpenCell. Number of protein pairs per category are shown above the bars.

**Figure S4.**
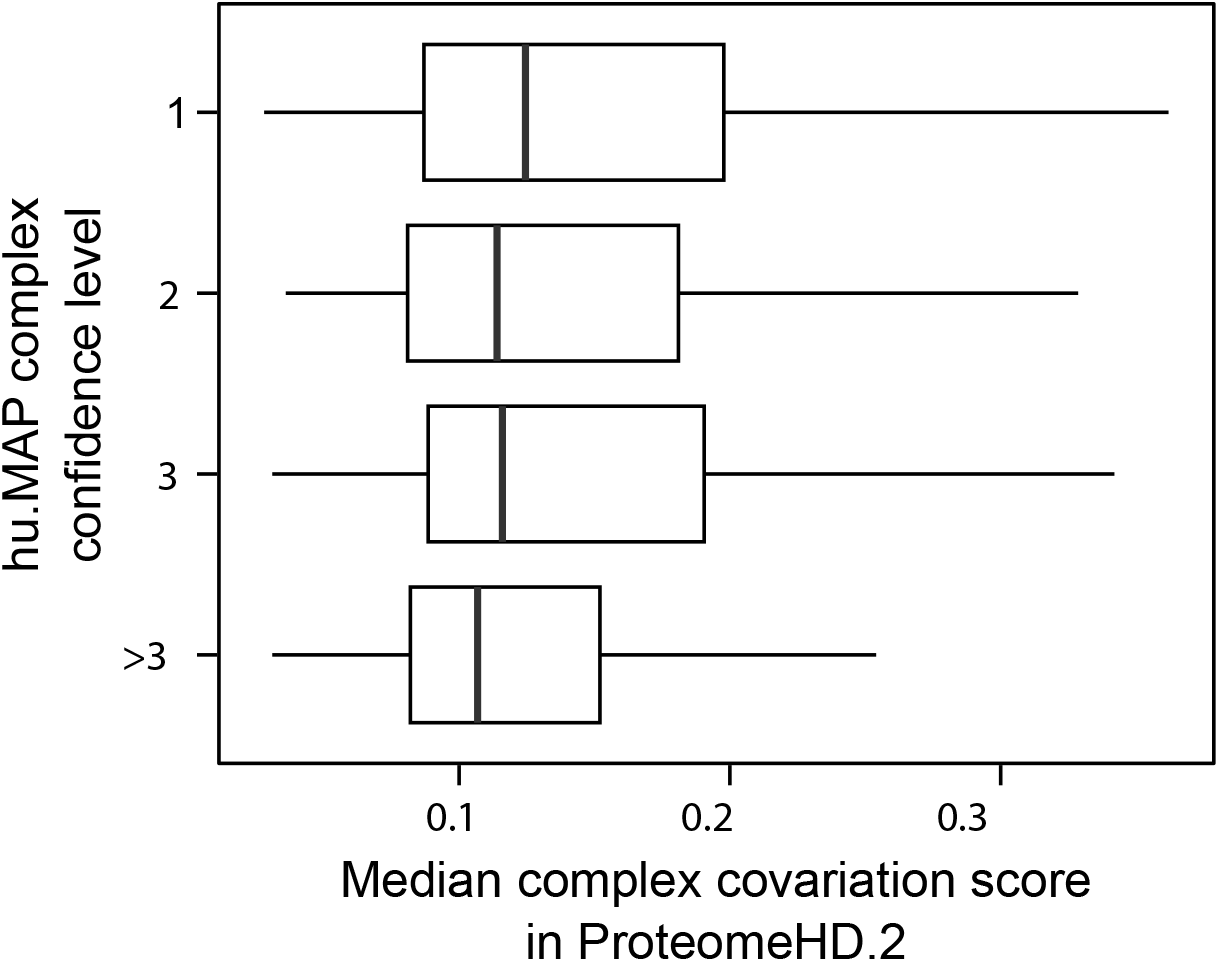
Median complex covariation does not strongly correlate with hu.MAP3.0 confidence levels. The median of protein pairwise covariation probability from ProteomeHD.2 for subunits belonging to the same hu.MAP3.0 complex was calculated, for complexes that had more than 50% coverage in ProteomeHD.2.

**Figure S5.**
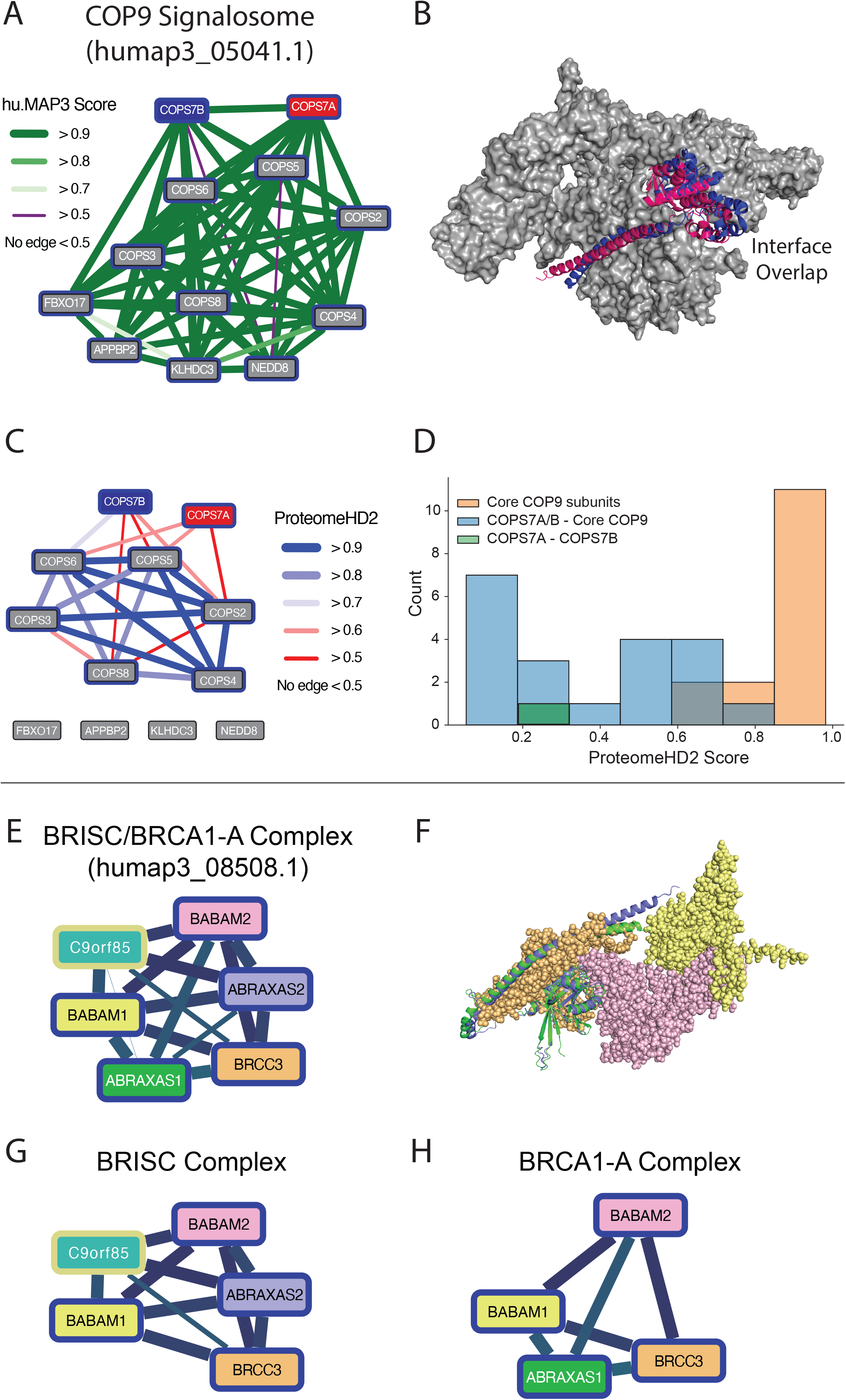
Example complexes with mutual exclusive subunits. (**A**) Co-complex interaction network of the COP9 Signalosome shows high interconnectivity of core subunits and peripheral subunits FBXO17, APPBP2, KLHDC3, and NEDD8. (**B**) Structure of COP9 Signalosome core subunits show overlapping interfaces of COPS7A and COPS7B with other core subunits (PDB id: 4D10, 6R6H) pointing to these subunits being mutually exclusive. (**C**) Protein co-expression network of COP9 Signalosome from ProteomeHD.2. Core subunits of the network are highly co-expressed while COPS7A and COPS7B are less co-expressed to core subunits. (**D**) Distributions of co-expressions between core COP9 subunits (orange), COPS7A or COPS7B and core COP9 subunits (blue), and the co-expression between COPS7A and COPS7B (green). (**E**) The BRISC/BRCA1-A Complex highlights the identification of two mutually exclusive proteins and the annotation of an uncharacterized protein. Blue border of protein subunits represents a known member of the complex; the yellow border represents an uncharacterized protein. Edge weights represent hu.MAP3.0 confidence score. (**F**) AlphaFold3 model of known subunits of complex. The interfaces of ABRAXAS1 and ABRAXAS2 with BRCC3 overlap and therefore are determined to be mutually exclusive subunits. (**G-H**) BRISC/BRCA1-A Complex split into two complexes which appropriately separates mutually exclusive subunits and places the uncharacterized protein, C9ORF85, with the BRISC complex.

**Figure S6:**
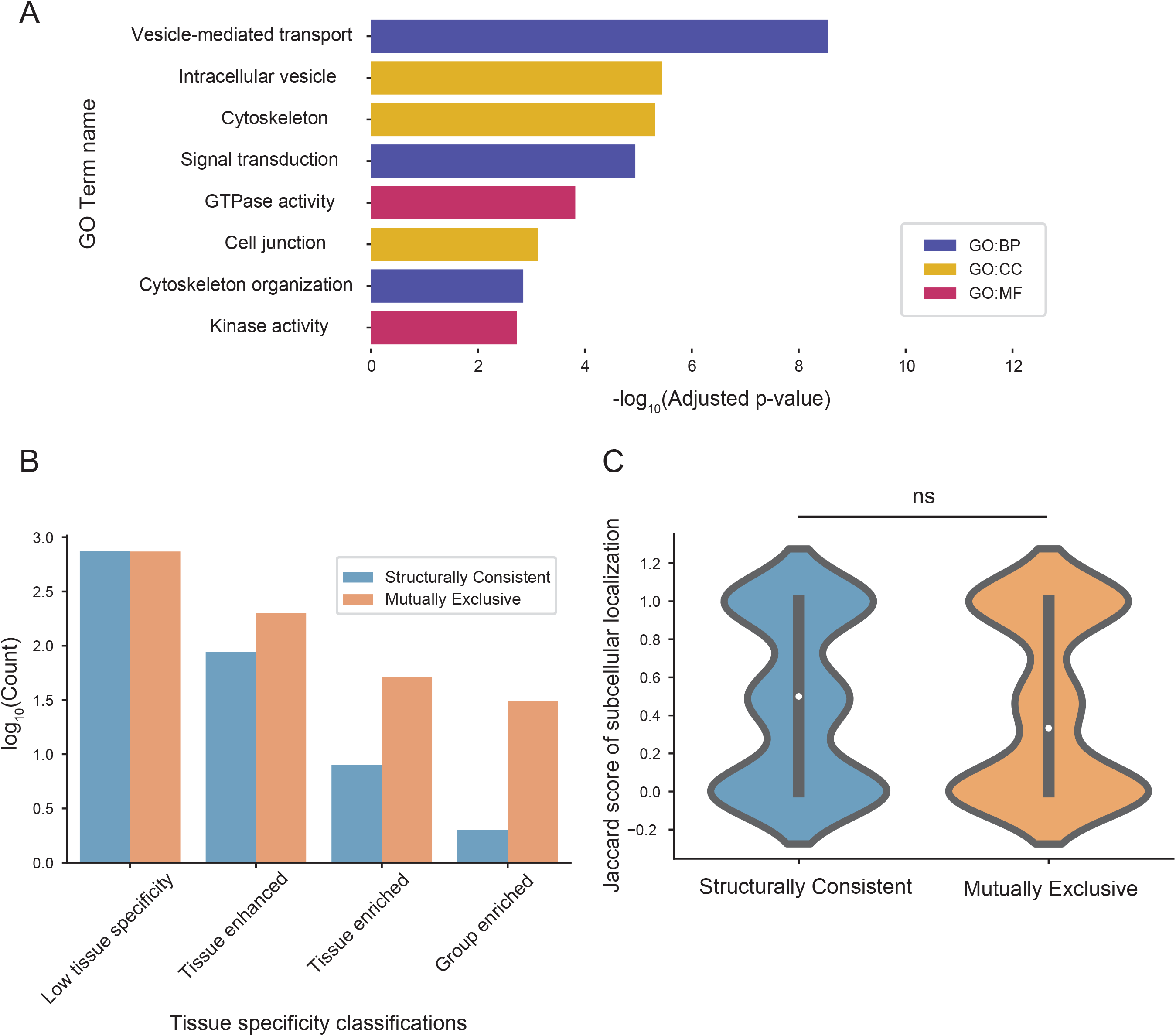
Characterization of mutually exclusive interactions. (**A**) Barplot displaying gene ontology annotation enrichments for molecular function (MF), biological process (BP), and cellular compartment (CC) for identified mutually exclusive proteins. (**B**) Barplot showing the count (log_10_ transformed) of proteins from mutually exclusive or structurally consistent protein pairs in different Human Protein Atlas tissue expression classifications. (**C**) Violin plot showing the distribution of Jaccard similarity of subcellular localizations between proteins in mutually exclusive and structurally consistent protein pairs (not significant; violin shape indicates density of data points with wider sections indicating higher concentration of values, the edges of the inner box indicate 25th and 75th percentile with inner dot indicating the median, the whiskers extend to minimum and maximum values).

**Figure S7:**
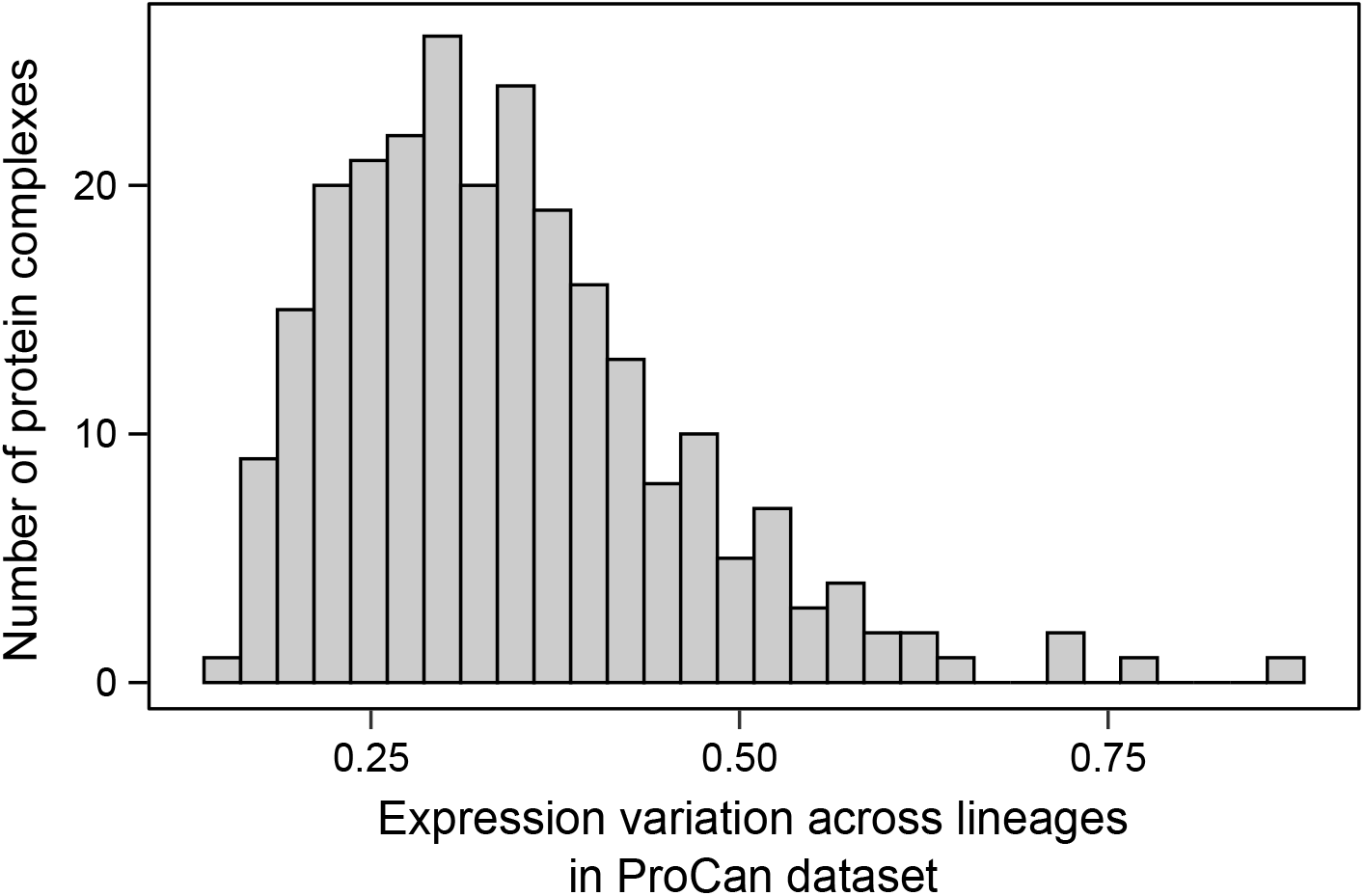
**Complex abundance levels vary between cancer lineages** Relative protein expression across cancer lineages was averaged to hu.MAP3.0 complex-level to identify complexes that differ in abundance between lineages, as quantified by the standard deviation of the complex-level z-score. Subunits with more than 80% missing values in the ProCAN dataset were excluded from downstream analysis.

**Figure S8:**
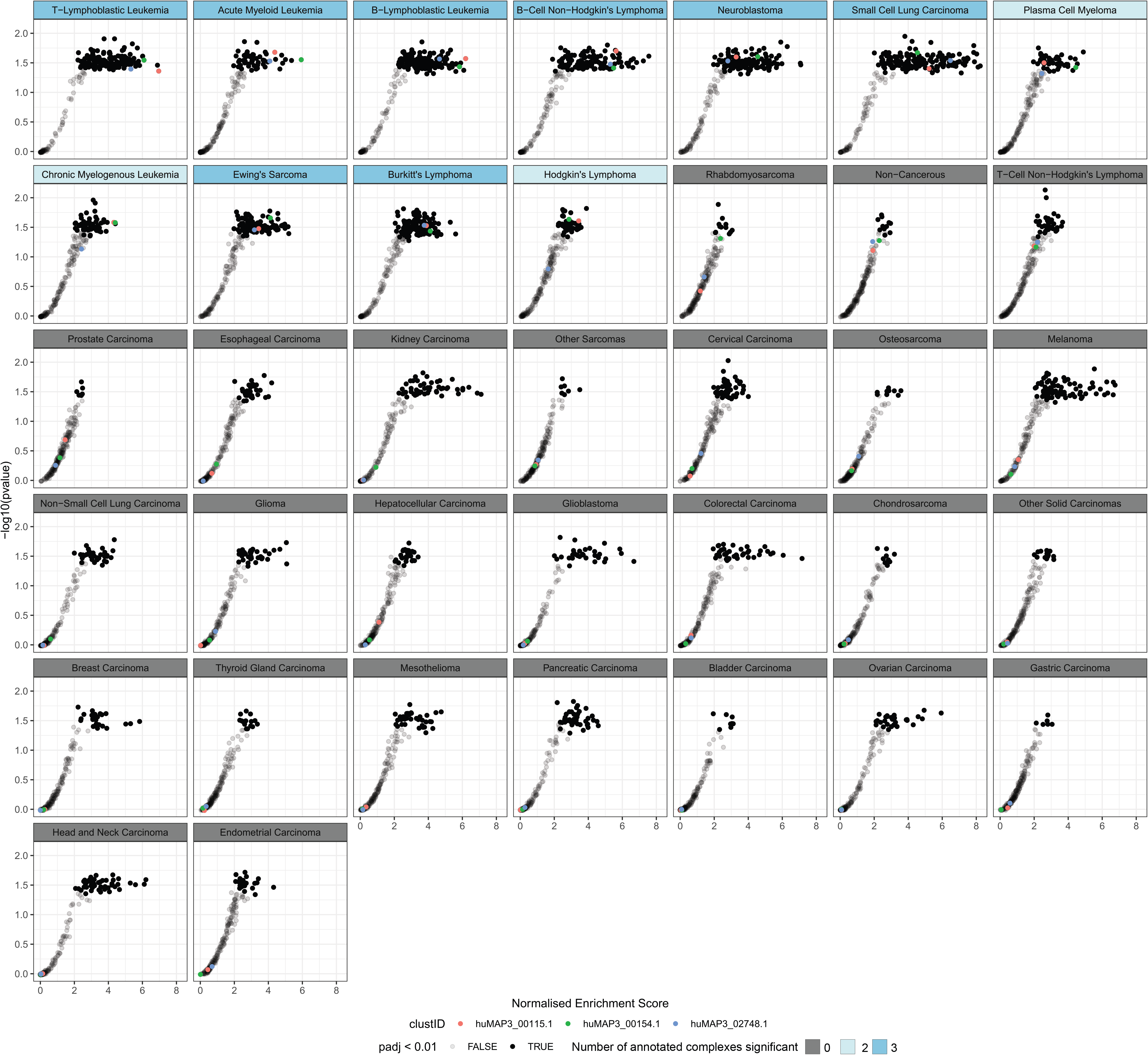
**Statistical enrichment of hu.MAP3.0 complex abundance across cancer lineages** An overrepresentation test was performed to identify complexes that are overexpressed in certain cancer lineages. Each point represents a hu.MAP3.0 complex, with its relative lineage-enrichment score on the x-axis and the overrepresentation significance on the y-axis. Complexes with FDR-adjusted p-values below 0.01 are shown in black and selected complexes are annotated. Lineages with significant overexpression of all three of these selected complexes are shown in dark blue, and those with two complexes reaching significance in light blue. Protein subunits with more than 80% missing values in the cancer lineage dataset were excluded from this analysis.

## Notes

### Competing Interest Statement

The authors have declared no competing interest.

